# Preclinical evaluation of the efficacy of α−Difluoromethylornithine and Sulindac against SARS-CoV-2 infection

**DOI:** 10.1101/2025.02.26.640194

**Authors:** Natalia A. Ignatenko, Hien Trinh, April M. Wagner, Eugene W. Gerner, Christian Bime, Chiu-Hsieh Hsu, David G. Besselsen

**Affiliations:** Department of Cellular and Molecular Medicine, University of Arizona, Tucson, AZ; Valley Fiver Center for Excellence, University of Arizona, Tucson, AZ; University Animal Care, University of Arizona, Tucson, AZ; Department of Medicine, Division of Pulmonary and Critical Care Medicine, University of Arizona College of Medicine, Tucson, AZ; Mel and Enid Zuckerman College of Public Health, University of Arizona, Tucson, AZ

**Author notes:** Correspondence to: Natalia A. Ignatenko, Department of Cellular and Molecular Medicine, University of Arizona Cancer Center, 1515 N. Campbell Ave., P.O. Box 245024, Tucson, AZ 85724-5024.

**Keywords:** SARS-CoV-2, COVID-19 mouse model, polyamines, DFMO, Sulindac, antiviral drugs

## Abstract

Despite numerous research efforts and several effective vaccines and therapies developed against COronaVIrus Disease 2019 (COVID-19), drug repurposing remains an attractive alternative to identify new treatments for SARS-CoV-2 virus variants and other viral infections that may emerge in the future. Cellular polyamines support viral propagation and tumor growth. Here we tested the antiviral activity of an irreversible inhibitor of polyamine biosynthesis, α-difluoromethylornithine (DFMO) and a non-steroidal anti-inflammatory drug (NSAID) Sulindac, which have been previously evaluated for colon cancer chemoprevention. The drugs were tested as single agents and in combination in human Calu-3 lung adenocarcinoma and Caco-2 colon adenocarcinoma cell lines and the *K18-hACE2* transgenic mouse model of severe COVID-19. DFMO/Sulindac combination significantly suppressed SARS-CoV-2 N1 Nucleocapsid mRNA and *ACE2* mRNA levels in the infected human cell lines by interacting synergistically when cells were pretreated with drugs and additively when treatment was applied to the infected cells. The antiviral activity of DFMO and Sulindac was tested *in vivo* as prophylaxis (drug supplementation at the doses equivalent to the human chemoprevention trial started 7 days before infection) or as treatment (drug supplementation started 24 hours post-infection). Prophylaxis with DFMO and Sulindac as single agents significantly increased survival rates in the young male mice (p=0.01, and p=0.027, respectively), and the combination was effective in the aged male mice (p=0.042). Young female mice benefited the most from the prophylaxis with Sulindac alone (p=0.001) and DFMO/Sulindac combination (p=0.018), while aged female mice did not benefit significantly from any interventions. The treatment regime was ineffective in suppressing SARS-CoV-2 infection in *K18-hACE2* mice. Overall, animal studies demonstrated the protective age- and sex-dependent antiviral efficacy of DFMO and Sulindac against SARS-CoV-2.

## Introduction

Many RNA viruses, such as influenza, Chikungunya (ChK), Zika, Ebola, and coronaviruses, present a significant danger to the human population. Coronaviruses, particularly, can impose major economic loss worldwide, which has become especially evident when the novel, highly pathogenic Severe Acute Respiratory Syndrome CoronaVirus 2 (SARS-CoV-2) virus emerged. The SARS-CoV-2 virus was highly pathogenic and caused a severe lower respiratory tract infection with acute respiratory distress and extrapulmonary organ dysfunction in infected vulnerable individuals, which resulted in a loss of more than 7 million human lives worldwide (1).

Detailed analysis of COVID-19 cases revealed significantly higher vulnerability of the older population (75 years and older) with underlying health conditions, such as diabetes, lung disease, cancer, immunodeficiency, asthma, kidney disease, and gastrointestinal disease. Additionally, males have been more susceptible to SARS-CoV-2 with a higher rate of death (2). Even though effective vaccines and antiviral drugs specific for SARS-CoV-2 infection have been developed, there is a need to evaluate alternative prevention and treatment strategies to overcome potential resistance to existing anti-COVID-19 medications and to treat novel SARS-CoV-2 virus variants.

SARS-CoV-2 belongs to a family of single-stranded positive, enveloped RNA viruses, which cause respiratory, enteric, and systemic infections in animals and humans. SARS-CoV-2 infection starts with the binding of a virus to the transmembrane receptor angiotensin-converting enzyme 2 (ACE2) on the host cells with subsequent Spike (S) protein priming by a serine protease TMPRSS2 (3). Virus particles enter cells via endocytosis and replicate in the host’s cytoplasm through the synthesis of a full-length negative strand RNA by replicase, which serves as a template for genomic RNA. The viral RNA is translated at the endoplasmic reticulum (ER) membrane, and viral particles are transported through the ER-Golgi secretory pathway to the cell surface, where the virus is released (4). Moreover, single-stranded SARS-CoV-2 RNA initiates a cellular immune response with marked elevation in inflammatory mediators (the “cytokine storm”), which destroys cell integrity and leads to a release of viral particles (5,6).

Understanding virus dependency on host cell resources for replication presents opportunities to suppress and even eliminate the infection. Previous gene ontology (GO) enrichment analysis of RNA viruses’ interactions with human host cells showed that RNA viruses target mostly cellular protein sets involved in gene expression, with a focus on RNA processing, splicing, localization, and transport within the host cell. DNA viruses mostly interact with human protein sets involved in cellular chromatin assembly, nucleosome organization, DNA packaging, and conformational changes (7).

Numerous *in vitro* experiments have demonstrated the importance of cellular polyamines for viral replication in both animals and humans (reviewed in (8,9). Polyamines putrescine, spermidine, and spermine are naturally occurring organic cations that are essential for the growth and development of both prokaryotic and eukaryotic cells (10,11). Polyamines are involved in diverse cellular processes, including control of gene expression at the transcriptional, post-transcriptional, and translational levels (12,13). Polyamines can also influence the expression of pro-inflammatory genes, such as cyclooxygenase 2 (COX2) (14).

Intracellular polyamine levels are tightly regulated, which indicates that these molecules are important for optimal cellular growth (15–17). Polyamines are synthesized through the action of two rate-limiting biosynthetic enzymes: ornithine decarboxylase (ODC1), which catalyzes the formation of putrescine from ornithine, and S-adenosylmethionine decarboxylase (AdoMetDC). AdoMetDC provides methyl groups for the synthesis of spermidine from putrescine with the action of spermidine synthase (SPS), and the synthesis of spermine from spermidine by spermine synthase (SMS). Important catabolic effectors maintain polyamine homeostasis as well. Spermidine/spermine N-acetyltransferase (SAT1) adds terminal acetyl groups to propyl amine moieties of spermidine and spermine, facilitating their export as acetylated polyamines. Spermine oxidase (SMOX) catalyzes the conversion of spermine into spermidine with the secondary production of hydrogen peroxide. Acetylpolyamine oxidase (PAOX) also converts acetylated spermidine and spermine to putrescine and spermidine, respectively. A key polyamine catabolic enzyme, SAT1 is highly inducible in response to a variety of stimuli, including elevated polyamine levels, synthetic polyamine analogs, toxins, hormones, cytokines, heat-shock, ischemia-reperfusion injuries, and other stresses (reviewed in (16)). Polyamines can also be imported into the cells from extracellular sources such as luminal bacteria and diet (18). A synthetic irreversible inhibitor of ODC1 enzyme activity, eflornithine (α-difluoromethylornithine, or DFMO) (19) has become a valuable tool in assessing the function of polyamines in the normal and diseased state.

There is ample evidence that viruses require polyamines throughout their life cycle for genome packaging, DNA-dependent RNA polymerase activity, genome replication, and viral protein translation (20,21). The polyamines spermidine and spermine were detected in the viral envelope of herpes simplex virus (HSV-1) and have been shown to contribute to stabilizing capsids through interaction with the negatively charged viral DNA genome (34). Investigations of the polyamine function of mammalian viruses through an *in vitro* genome synthesis assay or polyamine inhibition by DFMO revealed that many viruses, including. chikungunya virus (CHIKV), Zika virus (ZIKV) (20), hepatitis C virus (HCV) (22), herpes simplex virus (HSV-1) (23), and even Ebola virus (EBOV) (24), rely on host polyamines for viral translation. In addition, viruses can stimulate host cell polyamine production, as has been shown for human cytomegalovirus (HCMV), which induces host ODC1 activity and increases polyamine uptake in infected cells (25,26).

A more rigorous evaluation of drugs that target polyamine metabolism for antiviral indications is needed, specifically to ameliorate the severity of SARS-CoV-2 infection. Previously reported data provide abundant evidence that polyamines play vital roles in viral infection and that depletion of polyamine levels in the host cells can suppress viral propagation. DFMO can restrict the replication of numerous RNA viruses, such as flaviviruses dengue virus serotype 1 (DENV1), Japanese encephalitis virus (JEV) and yellow fever virus (YFV), poliovirus (PV), bunyavirus Rift Valley fever virus (RVFV), and rhabdovirus rabies virus (RABV). Importantly, DFMO treatment of Middle East respiratory syndrome coronavirus (MERS-CoV)-infected Vero cells resulted in a 30-fold reduction in viral titer (20). Inhibition of ODC1 enzyme activity by DFMO also resulted in suppression of the translation of viral transcripts due to a decrease in the cellular spermidine level, which is required to hypusinate cellular translation factor eIF5A (24). These findings indicate that DFMO antiviral activity is multimodal and virus-specific and could be highly effective against the SARS-CoV-2 virus. Stimulation of polyamine excretion using synthetic polyamine analogs, which induce transcription of SAT1 catabolic enzyme, also has been shown to limit the replication of RNA viruses (20).

The polyamine depletion strategy based on a combination of inhibition of polyamine biosynthesis by DFMO and activation of polyamine excretion using the non-steroidal anti-inflammatory drug (NSAID) Sulindac has been shown to cause a depletion of intracellular polyamine levels and suppression of cell proliferation (27–29). DFMO and Sulindac as a combination treatment, have been utilized to prevent colorectal adenomas in patients with prior colon polyps and were proven safe and remarkably effective in preclinical in vivo studies and several clinical trials (30–32).

The goal of this study was to evaluate the efficacy of DFMO and Sulindac against SARS-CoV-2 infection in cell lines and a transgenic *K18-hACE2* mouse model of severe COVID-19 disease (33). We found that DFMO and Sulindac combination treatment significantly suppress SARS-CoV-2 virus propagation and *ACE2* mRNA levels in the infected cell lines. The *in vivo* experiments showed sex- and age-dependent protective antiviral activity of DFMO and Sulindac, when tested as a prophylaxis regime. Increased survival rates were observed in the young male mice receiving DFMO or Sulindac as single agents, and the DFMO/Sulindac combination was effective in the aged male mice. Young female mice demonstrated increased survival rates on the prophylaxis regime with Sulindac alone and DFMO/Sulindac combination, while aged female mice did not benefit significantly from any interventions.

## Materials and methods

### Materials

All cell culture reagents used in this study were purchased from Thermo Fisher Scientific, Inc. (Waltham, MA) and Sigma-Aldrich (St. Louis, MO). Difluoromethylornithine hydrochloride hydrate (DFMO, product number D193) was purchased from MilliporeSigma (Burlington, MA). Sulindac (Cat. # 10004386) was purchased from Cayman Chemical (Ann Arbor, MI). DFMO was diluted in Dulbecco’s Modified Eagle medium (DMEM) Stock solution of Sulindac for cell culture experiments was prepared in 100% Dimethylsulfoxide (DMSO) and diluted with DMEM.

### Cell lines

Vero rhesus monkey kidney cell line (ATCC-CCL-81, kindly provided by Dr. Janko Nikolich), Calu-3 lung adenocarcinoma cell line (ATCC-HTB-55), and Caco-2 colon adenocarcinoma cell line (ATCC-HTB-37) were maintained in DMEM (4.5 g/L glucose, L-glutamine, w/o sodium pyruvate, supplemented with 10% FBS and 1% penicillin/streptomycin). Cells were incubated at 37°C in 5% CO_2_.

### SARS-CoV-2 virus

All experiments involving SARS-CoV-2 were performed in compliance with the University of Arizona guidance for BSL-3 work. SARS-CoV-2 strain USA_WA1/2020 was obtained from Dr. Natalie Thornbutg through the University of Texas Medical Branch (UTMB) World Reference Center for Emerging Viruses and Arboviruses (WRCEVA). Virus was propagated as previously described (34). Briefly, 80-90% confluent Vero cells were infected with an approximate multiplicity of infection (MOI) of 0.01, incubated for 48 hours, and supernatant was harvested and clarified. Virus titer was determined by plaque assay.

### SARS-CoV-2 virus inactivation

The SARS-CoV-2 virus was inactivated using lysis buffers compatible with sample processing for downstream applications, with 20x Lysis RNase Inhibitor buffer (2% Igepal CA-630 (NP40), 3mg/ml Poly-vinylsulfonic acid (PVSA) solution in PBS) added to each sample for a final concentration of 1x. A 0.1% concentration of NP-40 in has been shown to completely inactivate SARS-CoV2 virions in lysed cells, and PVSA at a final concentration of 150 μg/ml has been shown to inhibit RNases (35,36). In cell culture experiments, after media was removed and infected cells were processed for viral RNA isolation, the virus was inactivated using lysis buffer RAV1 (NucleoSpin RNA Virus kit, Macherey-Nagel GmbH&Co, Dueren, Germany).

### Drug synergy analysis

Cell lines were seeded in 96 well plates at a concentration of 2×10^4^ cells per well (Vero, Caco-2 cell lines) or 3×10^4^ cells per well (Calu-3 cell line) in 50 μl media volume per well. For the prophylaxis regime, 24 hours after subculture cell culture media was replaced with media containing drugs, and cells were cultured for an additional 48 hours before exposure to the virus. For the treatment regime, cells were incubated in drug-free media for 48 hours before being infected. Under both regimes, cells were infected with the virus at an MOI of 0.05 for one hour, and then the virus-containing media was replaced with new media containing DFMO, Sulindac, or a combination of DFMO and Sulindac (DFMO/Sul combo). The infected control and treated cultures were harvested at 48 hours post-infection.

The tested drug concentrations ranged from 0.15 mM to 5 mM for DFMO and from 0.1 mM to 1.6 mM for Sulindac and were based on the previously reported concentrations of DFMO (150 to 500 μM), which resulted in ≥ 50% reduction in replication for multiple viruses from different viral families, including MERS-CoV (20). Variable Sulindac concentrations were selected based on the previously reported ability of sulindac and its metabolites to inhibit proliferation or induce apoptosis in different lung and colon cancer cells at 50-400 μM (29,37,38). These concentrations also span the serum concentrations achieved by a single oral daily dosing of DFMO and Sulindac used in patients’ therapeutic regimes.

The infected cell lines were treated with the tested drugs for 72h in duplicate. The virus was neutralized, and cells lysed with 20x Lysis RNase Inhibitor buffer added to a final concentration of 1x in a total volume of 100 μl. Contents of the wells were mixed, plates were frozen, and 5 μl of the lysed cultures were used as a template for measuring the level of viral nucleocapsid gene (N1) transcript by quantitative RT PCR (qRT-PCR). The percent of N1 inhibition by DFMO/Sulindac combo relative to the untreated control was calculated and further analyzed using a free web application SynergyFinder (https://synergyfinder.fimm.fi) (39). The Zero Interaction Potency (ZIP) model (40) was utilized to evaluate the potential synergy of drug combinations. The ZIP model interprets results based on the calculated summary score with a statistical significance (p-value). The mean ZIP value increases as the synergy between drugs increases. Specifically, a score less than -10 indicates antagonistic drug interaction, a score between -10 and 10 indicates the drugs have an additive effect, and a score >10 indicates synergistic effect.

### Cell culture experiments

The antiviral activity of the tested drugs was evaluated in the Calu-3 cell line. Cells were seeded in 6-well or 24-well plates at 0.3 ×10^6^ cells/well or 0.5 ×10^5^ cells/well, respectively, in DMEM. Twenty-four hours after subculture the cells were infected with SARS-CoV-2 at a MOI of 0.05. After 1 h, the virus-containing media was replaced with media containing the tested compounds at the following concentrations: 1 mM DFMO, 300 μM Sulindac, or DFMO/Sul combo (1 mM and 300 μM), while the infected untreated cells were incubated with vehicle control (DMSO). Treatments of the infected cells were done in triplicates. At 72 h post-infection, the virus was inactivated and cells were lysed as described above. Samples were frozen until analyzed by qRT-PCR to quantify the drug effects on a viral load and host cell gene expression.

### Plaque Assay

The SARS-CoV-2 plaque-forming assay was done using a liquid overlay and fixation-staining method according to the previously described protocols (34,41). Briefly, Vero cells were plated in 24-well plates and infected with 10-fold serial dilutions of virus stock or clarified lung supernatant once cells reached 90% cell confluency. Plates were incubated for 1-2 hours at 37^0^C in 5% CO_2_. Cell culture media containing methylcellulose (1% w/v) was added to each well and plates were incubated for 72 hours. Media was discarded and each well was filled with 10% formalin and left for 30 minutes to inactivate any virus and fix cells. Plates were washed with tap water and then the bottom of each well was filled with 0.9% Crystal Violet (w/v) in 40% ethanol solution to stain cells. The plates were washed and dried and the plaques were counted to calculate titer in PFU/ml.

### RNA isolation

Viral RNA from the lysed cells was isolated for host cell expression analysis using the NucleoSpin RNA Virus kit (Cat.#740956, Macherey-Nagel GmbH&Co) according to the manufacturer’s protocol. Total RNA extraction was performed using the Qiagen RNeasyPlus Kit (Cat. # 74136, Qiagen GmbH, Hilden, Germany) according to the manufacturer’s protocol. RNA purity was assessed using a Nanodrop spectrophotometer (ND-2000; Thermo Fisher Scientific, Inc.).

### Quantitative reverse-transcription PCR (qRT-PCR)

Five microliters of each lysed cell culture was used directly as a qRT PCR template to measure the expression of the SARS-CoV-2 viral gene, encoding N1 Nucleocapsid protein. Absolute quantification of N1 was performed using SARS-CoV-2-specific in-vitro-transcribed N1 RNA standard (2019-nCoV_N_Positive Control) and primers (42,43) using QuantaBio qScript XLT One-Step qRT-PCR ToughMix (Cat.# 89236-676, VWR International, Randor, PA). TaqMan® SARS-CoV-2 assays were purchased to detect the Spike glycoprotein S1 subunit 1 (Cat. 3 4221182, Assay ID # V107918636-S1) and human RNase P (Cat. 3 4331182, Assay ID # Hs04930436-g1, Thermo Fisher Scientific, Inc.), which was used for an extraction and internal amplification quality control for cell culture experiments. For N1 RT qPCR analysis in animal experiments, 5 μl of lung tissue homogenate was used as a template and the N1 transcript number was normalized to the expression level of the mouse glyceraldehyde-3-phosphate dehydrogenase (GAPDH) gene (Cat # 4331182, Assay ID # Mm99999915_g1) in the same sample. For the host gene expression analysis, reverse transcription was completed using the Applied Biosystems High-Capacity cDNA Reverse Transcription Kit (Cat. # 4368814, Applied Biosystems, Foster City, CA). Total RNA (2 μg) was transcribed into cDNA in a 100 μl reaction using random hexamers under thermal conditions recommended by the manufacturer. qRT-PCR was performed using human TaqMan® assays specific for ornithine decarboxylase, *ODC1* (Cat. # 4331182, Assay ID # (Hs00159739_m1), spermine oxidase *SMOX* (Assay ID # Hs00602494_m1), spermidine/spermine acetyltransferase, *SAT1* (Assay ID # Hs00161511_m1), *ACE2* (Assay ID# Hs01085333_m1), and β2-microglobin (β2M, FAM (Hs99999907_m1) which was used as an endogenous reference for the host gene transcript level. Total RNA (0.2 µg) was reversed transcribed into cDNA in a 20 µL reaction with random hexamers under thermal conditions recommended by the protocol. Real-time PCR amplification was performed using the QuantStudio 3.0 Real-Time PCR system (Applied Biosystems, Life Technologies, Inc.), under the universal thermal cycling conditions recommended by the Assay-on-Demand products protocol. Negative controls without templates were included in each plate to monitor potential PCR contamination. The expression of genes was tested in triplicate and each reaction was run in duplicate. The comparative *C_t_* method was used to determine the relative expression level of each target gene. The *C_t_* value of each target gene was normalized by the endogenous reference. The fold change in expression of each target gene was calculated via the equation 2^-ΔΔ^*^Ct^* where ΔΔ*C_t_* = (*C_t_*_(target)_ – *C_t_*_(endogenous control)_.)_treatment_-( (*C_t_*_(target)_ – *C_t_*_(endogenous control)_)_control_.

### Animal experiments

Six-week-old *C57BL/6J K18-hACE2* transgenic mice of both sexes were purchased from Jackson Labs (https://www.jax.org/strain/034860). *K18-hACE2* mice express human angiotensin-converting enzyme 2 (ACE2) under the control of the promoter and first intron of the human cytokeratin 18(K18) gene in epithelia, including airway epithelia, and recapitulates severe coronavirus disease (33). One set of the animals was used in experiments at 6 weeks old (young mice groups), and another set was maintained at the facility and used in experiments upon reaching 58 weeks of age (aged mice groups). All mice were maintained in the University of Arizona’s Animal Care Facility in accordance with The University of Arizona Institutional Animal Care and Use Committee (IACUC). Animals were housed in groups of five in individually ventilated microisolator cages under fluorescent lighting on a 14:10 light:dark cycle. Irradiated feed (Teklad Global diet, Cat. # 2919, Inotiv, Indianapolis, IN) and hyperchlorinated reverse osmosis drinking water were available *ad libitum.* The animals were fed a defined synthetic diet AIN-93G (Cat. 3 TD.94045, Inotiv) for 7 days prior to the start of experiments.

#### Administration of compounds

For the prophylaxis regime in 6-week-old and 58-week-old mice and the treatment regime in 58-week-old mice, DFMO at a dose of 835 ppm and Sulindac at a dose of 167 ppm were administered in custom-prepared AIN-93G diets. These dietary preparations of DFMO and Sulindac have been successfully used in murine cancer models and are based on the recommended human drug doses (30,44,45). The treatment regime in 6-week-old mice was done by intragastric gavage (IG) starting at 24 h after mice were infected with 1000 PFU of SARS-CoV-2. The drugs were administered by IG daily in 0.1 ml volume per mouse/day at doses equivalent to the doses of DFMO and Sulindac in the clinical trial for the prevention of sporadic colorectal adenomas (45). The aqueous solutions of DFMO at 154.17 mg/kg (mg of compound per kg of body weight) in water pH 11, Sulindac at 30.8 mg/kg in 0.25 M sodium bicarbonate, DFMO/Sulindac combo in 0.25 M sodium bicarbonate, or 0.25 M sodium bicarbonate as a vehicle control. The gavage needles were pre-coated with 1g/ml of sucrose before gavage to improve the swallowing reflex and decrease the time for passing the needle into the esophagus. Reverse osmosis drinking water was available *ad libitum* for the duration of the experiments.

#### Animal infection and monitoring

Mice were inoculated with SARS-CoV-2 virus intranasally at a dose of 1000 plaque-forming units (PFU) immediately after brief sedation with inhaled isoflurane. The standardized SARS-CoV-2 dose of 1000 PFU was selected for mouse infections based on a 20% survival rate at 14 dpi in 6-week-old *K18-hACE2* mice. Non-invasive body weight and clinical symptom scoring were performed daily. Specifically, infected animals were observed daily for signs of body weight loss, rapid breathing, hunched posture, lethargy, inactivity, and/or rough hair coat. The following clinical scoring was used for weight: 0, no weight loss; 1, <10 % weight loss; 2, 10-19% weight loss; 3, >20% weight loss; for breathing: 0, normal; 1, mild tachypnea (mild increase in RR); 2, severe tachypnea (marked increase in RR); 3, dyspnea (deep, labored breathing), and overall status: 0, healthy; 1, less than normal activity; 2, inactive; 3, no movement; 4, no response to stimuli; and 5, dead. Mice were considered moribund and euthanized if they exhibited a >4 ° C drop in body temperature, >20% body weight loss, severe dyspnea, inability to ambulate, inability to access food/water, paralysis, or non-responsiveness to stimulation.

#### Sample collection and processing

Animals were euthanized 10-14 days post-infection using isoflurane. Blood and 80-100 mg of lung tissue were collected. Blood was added to EDTA, centrifuged and 25 μl of plasma was treated with NP-40 (to inactivate virus) and frozen at 80° C until processed for polyamine and Sulindac metabolite analysis. Tissue was homogenized with glass beads in 500 μl PBS using a Mini-Beadbeater-96 (Biospec Products, Bartlesville, OK) at a maximum speed of 2400 rpm/s for 2 min following centrifugation at 5000 rpm for 10 min at room temperature. A 100 μl aliquot of supernatant was removed and treated with 20xLysis RNase Inhibitor to inactivate virus as stated previously. The rest of the supernatant was evaluated by plaque assay (and stored frozen).

### Polyamine and Sulindac metabolites analysis

Ten microliters of plasma samples were assayed for polyamine content by reverse-phase high-performance liquid chromatography (HPLC) with 1,7-diaminoheptane as an internal standard (46). Amines detectable by this method included putrescine, cadaverine, histamine, spermidine, spermine with a limit of detection of 5 pmol. Data are expressed as nmol polyamine per ml. The HPLC analysis of sulindac and its metabolites sulindac sulfide and sulindac sulfone was done in plasma samples diluted 1:50. The aliquots of diluted serum were mixed with the internal standard solution (100 ng/mL of indomethacin in acetonitrile) and processed according to the previously described method (47).

### Statistical analysis

Statistical analysis of viral and host gene expression in cell lines was done using ANOVA single-factor test. A p-value <0.05 was considered statistically significant. Kaplan-Meier estimation was performed to derive the median survival time by treatment in different experimental groups of animals. Cox regression was performed to derive the hazard ratio (vs. control) for each treatment group. The clinical scores of animals at the time of euthanasia were summarized using mean±SD by group. A one-way ANOVA test was performed to compare clinical scores between treatment groups. Polyamine content and Sulindac metabolite levels in plasma were summarized using mean±SD by the group. One-way ANOVA/two-sample t-test was performed to compare polyamine content and Sulindac metabolites between groups. P-value (vs. untreated control) by sex was derived from one-way ANOVA and the interaction p-value between treatment and sex was derived from two-way ANOVA with the interaction terms between Treatment and Sex indicators.

## Results

### Evaluation of an antiviral activity of DFMO and Sulindac *in vitro*

#### Drug synergy analysis

The Calu-3 and Caco-2 human cell lines represent the anatomical sites (pulmonary and intestinal) of SARS-CoV-2 infection in humans because they express ACE2 receptor and are receptive to SARS-CoV-2 infection (5,48,49). We used these cell lines to assess whether DFMO combined with Sulindac has a stronger antiviral activity against SARS-CoV-2 than a single agent. The results of DFMO and Sulindac drug interaction analysis showed that combinations of DFMO/Sul act synergistically to suppress SARS-CoV-2 N1 transcript levels in Calu-3 and Caco-2 cells in a prophylaxis setting (Calu-3 cells Mean ZIP is 79820372.69, p=2.75e-1, and Caco-2 cells Mean ZIP is 253.89, p=3.15e-02), and the drug exert an additive inhibitory effect in a treatment setting (Calu-3 cells Mean ZIP score is -3.46, p=2.62e-01, and Caco-2 cells Mean ZIP score is -6.68, p=3.75e-01) (**Fig. 1A, B).** It is worth noting that the half maximum inhibitory concentrations (IC50) of DFMO were lower in a prophylaxis setting than in a treatment setting. The Vero cell line, which is commonly used for propagation of SARS-CoV-2 and generation of viral stock, exhibited a synergistic inhibitory effect in a prophylaxis setting with a much lower ZIP Mean synergy score of 18.89 (p= 3.35e-02), and a similar additive inhibitory effect in a treatment setting (Mean ZIP score is -2.62, p=6.59e01) (**Fig. S1**). Overall, the drug synergy analysis suggested that the DFMO/Sul combination will be more potent for prophylaxis of the viral infection.

**Figure 1.**
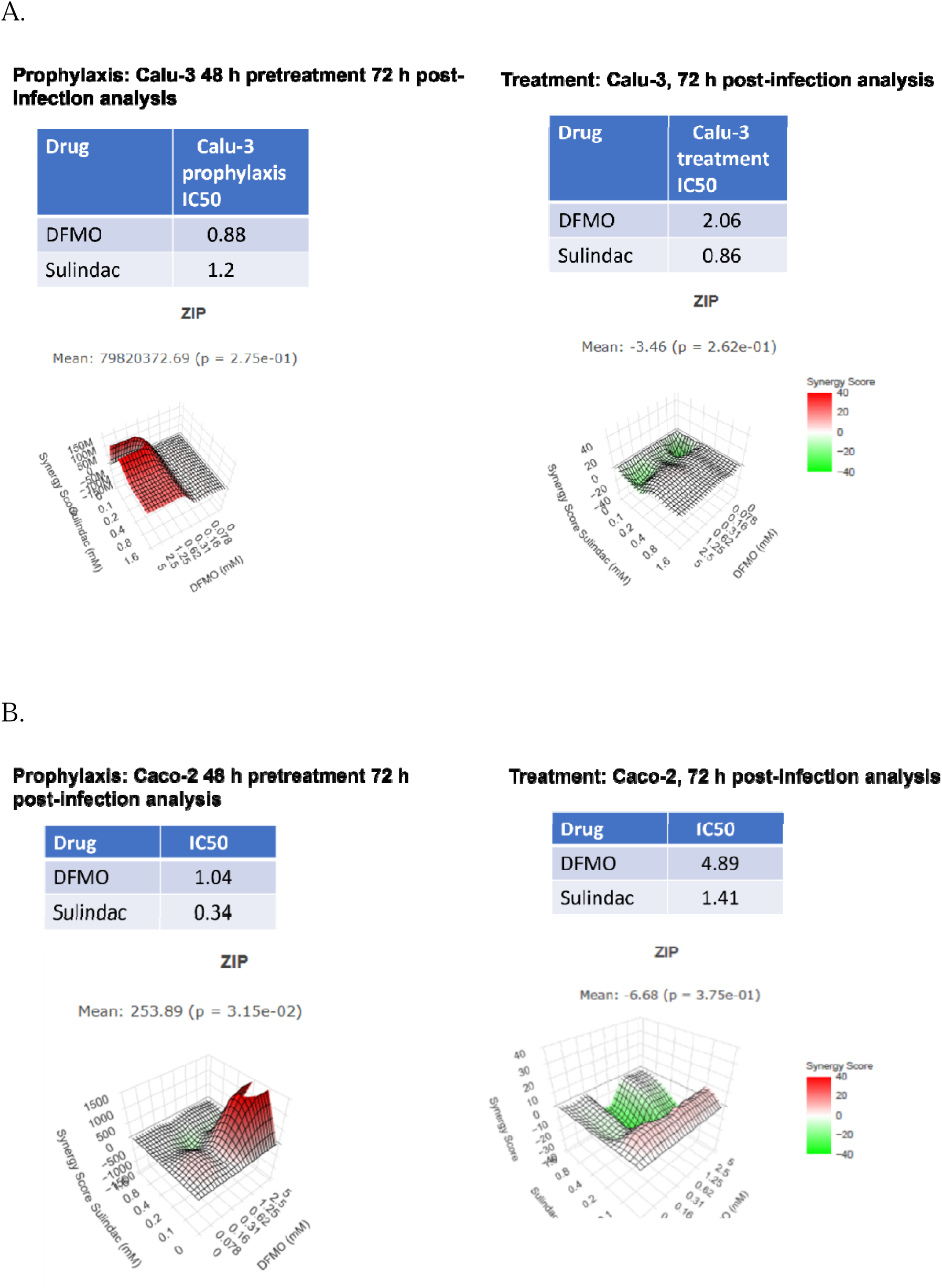
Results of SynergyFinder analysis of DFMO/Sul in Calu-3 and Caco-2 cell lines, infected with MOI 0.05. Cells were incubated with different concentrations of DFMO and Sulindac for 48 hours (Prophylaxis) before infection or drugs were added after cells were infected (treatment). N1 transcript level was measured by the qRT-PCR 72 hours post-infection.

#### Effect of DFMO and Sulindac on selected viral and host gene expression in infected Calu-3 adenocarcinoma cell line

SARS-CoV-2 viral replication relies on the host ACE2 receptor protein to infect and the host cell machinery to propagate within the cells. We measured N1 and Spike viral transcript levels in the conditional media and in the cell lysates of Calu-3 adenocarcinoma cells after cells were infected with SARS-CoV-2 virus at 0.05 MOI and incubated with DFMO and Sulindac either alone or in combination for 72 hours at concentrations below IC50 (1mM DFMO and 300 μM Sulindac). We observed a significant decrease in the copy number of SARS-CoV-2 N1 RNA in the conditioned media after treatment with Sulindac alone or in combination with DFMO (on average a 7-fold and a 10-fold, respectively) (P< 0.0001) **(Fig. 2A, left panel**). Analysis of N1 copy number in the infected cells lysates showed a 2-fold reduction by DFMO/Sul combo, but not the single agents (P<0.0001) **(Fig. 2A, right panel**). We noted a 2-fold increase in the N1 transcript copy number in the cell lysates of DFMO-only treated cultures compared to untreated cells or cells treated with Sulindac only or the DFMO/Sul combination (P<0.0001). This finding suggests that viral RNA processing may be suppressed in the host cells when DFMO inhibits polyamine synthesis. The level of SARS-CoV-2 Spike mRNA was reduced 5-fold in DFMO-treated cultures (P=0.02), and by more than 28-fold in Sulindac-treated cultures. A 14-fold reduction in Spike mRNA level was seen in Calu-3 cell lysates treated with DFMO/Sul combo (p<0.001) (**Fig. 2B**).

**Figure 2.**
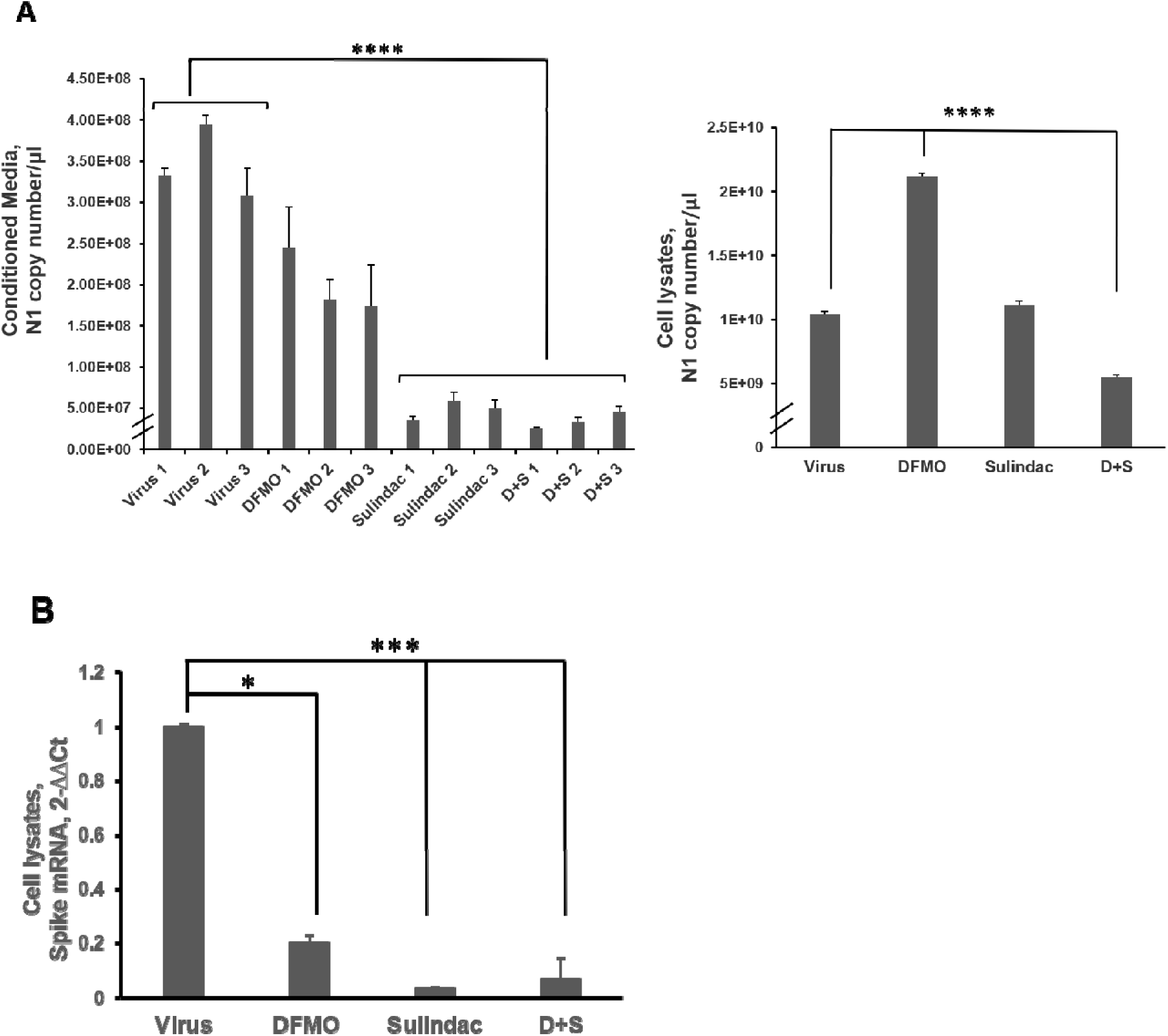
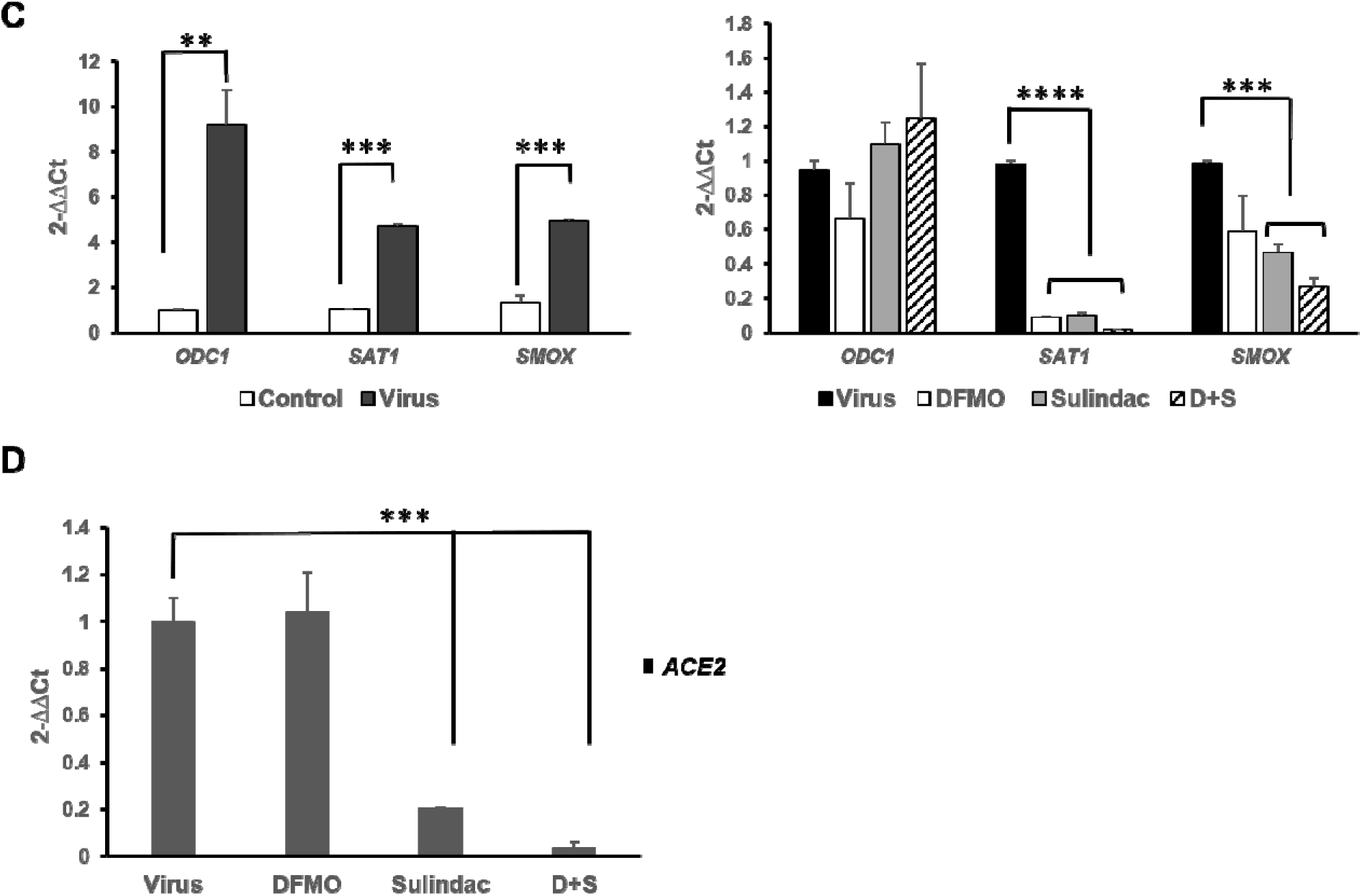
Analysis of the viral and host gene expression in Calu-3 cell line infected with SARS-CoV-2 at 0.05 MOI and treated with 1 mM DFMO, 300 μM Sulindac, or 1 mM DFMO/ 300 μM Sulindac combo at 72 h post-infection by qRT-PCR. **A. Left panel.** SARS-CoV-2 Nucleocapsid (N1) RNA copy number in the conditioned media of infected Calu-3 cells untreated (Virus,1-3), or treated with DFMO (DFMO,1-3), Sulindac (Sulindac, 1-3), and DFMO/Sulindac combo (D+S, 1-3) for 72 h post-infection. The N1 copy number was measured and the results from three independent experiments are shown. **Right panel**: N1 RNA copy number in cell lysates, infected untreated (Virus), DFMO-treated (DFMO), Sulindac-treated (Sulindac), and DFMO/Sulindac combo-treated (D+S). **B.** The transcript levels of Spike viral protein in cell lysates of the infected untreated and treated Calu-3 cells. Spike RNA level was normalized to the levels of the human RNase P in cells. Data were analyzed using ANOVA-single factor test *p =0.02, **p<0.01, ****p<0.0001. **C. Left panel.** The fold changes in expression of polyamine metabolic enzymes in Calu-3 cells infected with SARS-CoV-2 virus at MOI 0.05 (Virus). **Right panel.** Fold change in expression of polyamine metabolic genes in the infected Calu-3 cells due to treatment with DFMO, Sulindac, and DFMO/Sul combo for 72 hours (right panel). **D**. Fold change in expression of *ACE2* mRNA in Calu-3 cells infected untreated (Virus) and treated with compounds. Data were analyzed using ANOVA-single factor test *p <0.003. The results representative of three independent experiments are shown.

We also evaluated the mRNA levels of polyamine metabolic enzymes in uninfected (control) and infected Calu-3 cells treated with DFMO and Sulindac. Uninfected Calu-3 cells, incubated with the various concentrations of DFMO and Sulindac for 72 h, did not alter the *ODC1* and *SMOX* mRNA levels in a significant manner, except for the *ODC1* mRNA level in cells treated with DFMO/Sulindac combination. The *SAT1* level was significantly decreased in cells treated with DFMO at the concentration of 2.5 mM and 5 mM, as well as in cells treated with DFMO/Sul combination at 1 mM/100 μM and 2.5mM/200 μM (p<0.02) (**Fig. S2A**). The mRNA levels of polyamine metabolic enzymes *ODC1*, *SAT1,* and *SMOX*, were significantly elevated in SARS-CoV-2-infected cells compared to uninfected control cells (*ODC1* by more than 9-fold (p=0.03), *SAT1* and SMOX by 5-fold (p=0.0007 and p=0.008, respectively), which indicates that the virus induces polyamine metabolism (**Fig. 2C, left panel**). Treatment of infected Calu-3 cells with DFMO (1 mM) and Sulindac (300 μM) as single agents or in combination did not alter the *ODC1* mRNA level in the infected cells but significantly suppressed *SAT1* and SMOX genes expression (p=0.0007, p=0.002, respectively) (**Fig. 2C, right panel**). The *ACE2* mRNA level analysis in Calu-3 uninfected cells treated with the various concentrations of DFMO and Sulindac showed significant suppression of *ACE2* mRNA level in Sulindac alone and DFMO/Sul treated cultures, but not in DFMO-only treated cultures (p<0.001) (**Figure S2B**). Similarly, DFMO alone treatment did not affect the *ACE2* mRNA level in Calu-3 infected cells, but Sulindac alone and DFMO/Sul combination significantly reduced *ACE2* mRNA levels in the infected Calu-3 cells (by 5-fold and 28-fold, respectively). (**Fig. 2D,** p<0.003).

A analysis of viral gene expression was also performed in Caco-2 colon adenocarcinoma cell line treated with the various concentrations of DFMO and Sulindac (**Figure S3)**.

### Evaluation of DFMO/Sulindac combination in *K18-hACE2* mouse model of COVID-19

The initial assessment of the SARS-CoV-2 virus effect on plasma polyamine content in *K18-hACE2* mice was done by measuring the polyamine levels in animals infected with the SARS-CoV-2 virus at a dose of 1000 PFU at Day 7 post-infection. The polyamine levels in the infected young and aged animals of both sexes were compared with their level in matching uninfected animals (n=4 animals per group), which were maintained on an AIN-93G diet.

The infected 6-week-old male mice had more than 5-fold increase in the plasma polyamine levels, compared to the uninfected ones, mainly due to an increase in the level of putrescine (p<0.0001), which is synthesized from ornithine by ornithine decarboxylase (**Table 1A**). On the contrary, the infected aged male mice had a significantly lower level of polyamines than the uninfected aged mice due to the decrease in the concentration of putrescine (p<0.01) (**Table 1B**). These data indicate the sex-specific effect of SARS-CoV-2 viral infection with a significant increase in the plasma polyamine levels in the infected young male mice and a decrease in polyamine level in the infected aged male mice. The polyamine metabolism in the infected female mice was not significantly altered by the SARS-CoV-2 virus (**Table 1A**,**B**). Due to observed sex and age differences in plasma polyamine levels in the *K18-hACE2* mouse model, we continued the evaluation of DFMO and Sulindac antiviral efficacy within sex and age categories. Particularly, the antiviral activity of DFMO and Sulindac was assessed in young (6 weeks old) and aged (58 weeks old) infected *K18-hACE2* mice of both sexes after administration of the agents as a preventive measure (prophylaxis) and as a post-infection treatment (treatment) regime (**Fig. 3A,B**).

**Figure 3.**
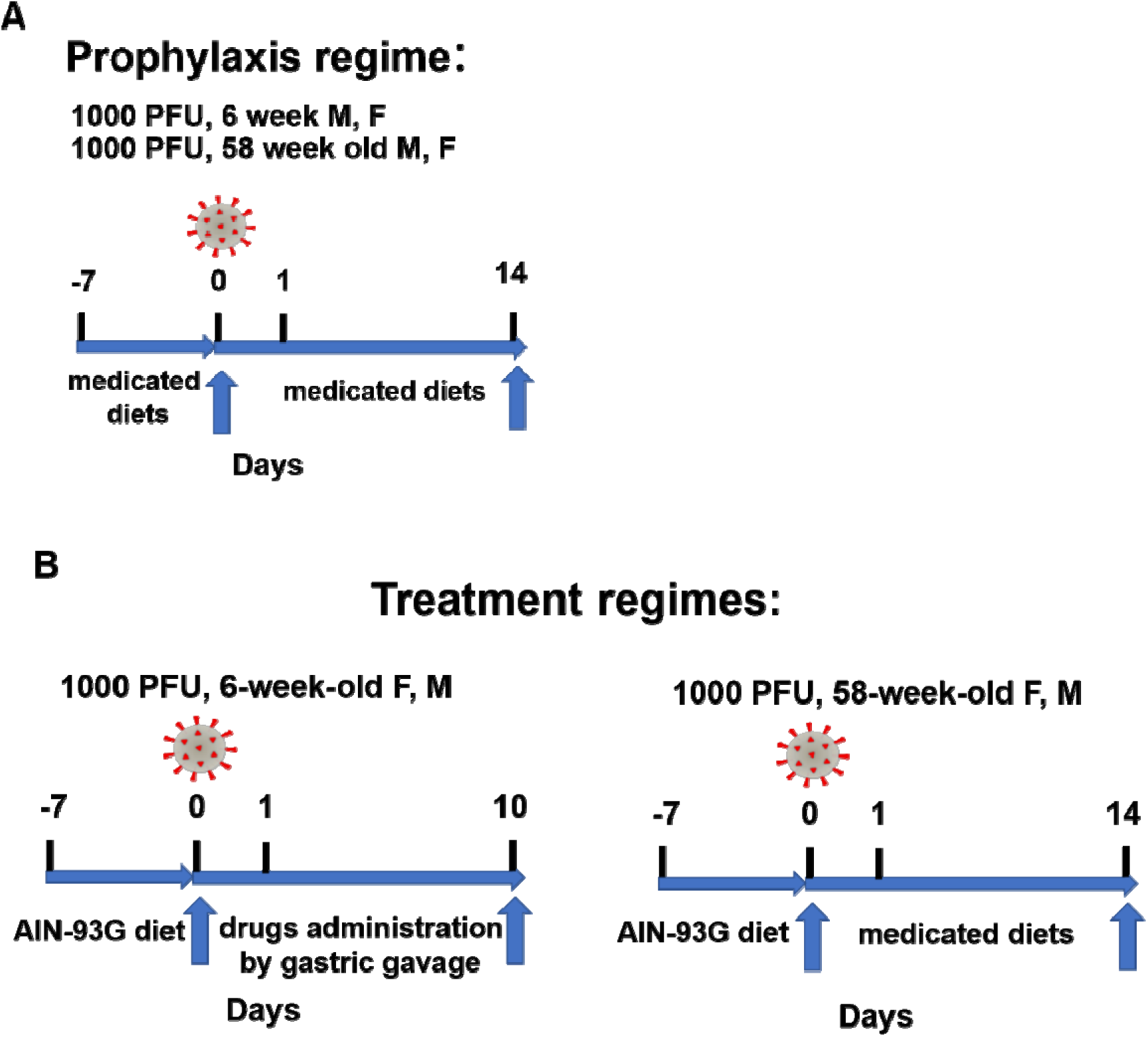
Experimental protocols with infection and treatment time points for animals placed on Prophylaxis regime (A) and Treatment regimes (B).

**Table 1.** A. Polyamine content in the plasma of control (not infected) and infected 6-week-old female and male *K18-hACE2* mice.

| **Polyamine content in 6-week-old mice (n=4/group) Mean ± SE (nmole/ml)** |  |  |  |  |  |  |  |  |
| --- | --- | --- | --- | --- | --- | --- | --- | --- |
| AIN-93G diet | Putrescine |  | Spermidine |  | Spermine |  | Total PA |  |
|  | female | male | female | male | female | male | female | male |
| Not infected | 13.64±1.12 | 0.59±0.08 | 4.10±2.16 | 2.37±0.37 | 0.21±0.18 | 0.28±0.05 | 18.33±3.20 | 3.24±0.45 |
| Infected, untreated | 18.06±1.14 | 18.27±1.74* | 3.78±1.42 | 1.47±0.31 | 0.21±0.17 | 0.03±0.01 | 22.26±2.56 | 19.78±2.05* |

\*p- value vs not infected <0.0001
n.d.-not detected

**Table 1.** B. Polyamine content in the plasma of control (not infected) and infected 58-week old female and male *K18-hACE2* mice.

n.d.-not detected B. Polyamine content in the plasma of control (not infected) and infected 58-week old female and male *K18-hACE2* mice.
| **Polyamine Content in 58-week-old mice (n=4/group). Mean ± SE (nmole/ml)** |  |  |  |  |  |  |  |  |
| --- | --- | --- | --- | --- | --- | --- | --- | --- |
| AIN-93G diet | Putrescine |  | Spermidine |  | Spermine |  | Total PA |  |
|  | female | male | female | male | female | male | female | male |
| Not infected | 14.16±0.89 | 23.05±2.47 | 1.48±0.28 | 1.17±0.18 | n.d. | 0.14±0.07 | 15.64±0.77 | 24.37±2.43 |
| Infected, untreated | 12.05±0.36 | 14.11±1.04* | 1.17±0.18 | 1.11±0.04 | n.d. | 0.04±0.02 | 13.47±0.23 | 15.26±1.10* |

\*p- value vs not infected <0.01
n.d.-not detected

### Prophylaxis regime in young mice

The 6-week-old, *K18-hACE2* female and male mice (5 per group) were placed on each of the four diets (AIN-93G, AIN-93G DFMO 835 ppm, AIN93G Sulindac 167 ppm, AIN-93G DFMO/ Sulindac combo) for 1 week before intranasal infection with 1000 PFU SARS-CoV-2, with continued drug supplementation as a medicated diet throughout the remainder of the experiment to a maximum 14 days post-infection (dpi) (**Fig.3A**).

#### Survival analysis

The survival rate of young untreated female mice was 0 at 14 dpi (mean survival 7 dpi). The survival rates did not improve in young female mice receiving DFMO alone diet (**Fig. 4A 6-week female mice)**. The young female mice treated with Sulindac alone had a statistically significant increase in survival (mean survival 11.6 dpi, p=0.001) with a statistically significant improvement in the clinical score (p=0.003). The young female mice treated with DFMO/Sulindac combo also showed a statistically significant increase in survival compared to the untreated mice (mean survival 7.8 dpi, p=0.018), even though no improvement in the clinical score was noted in this group.

**Figure 4.**
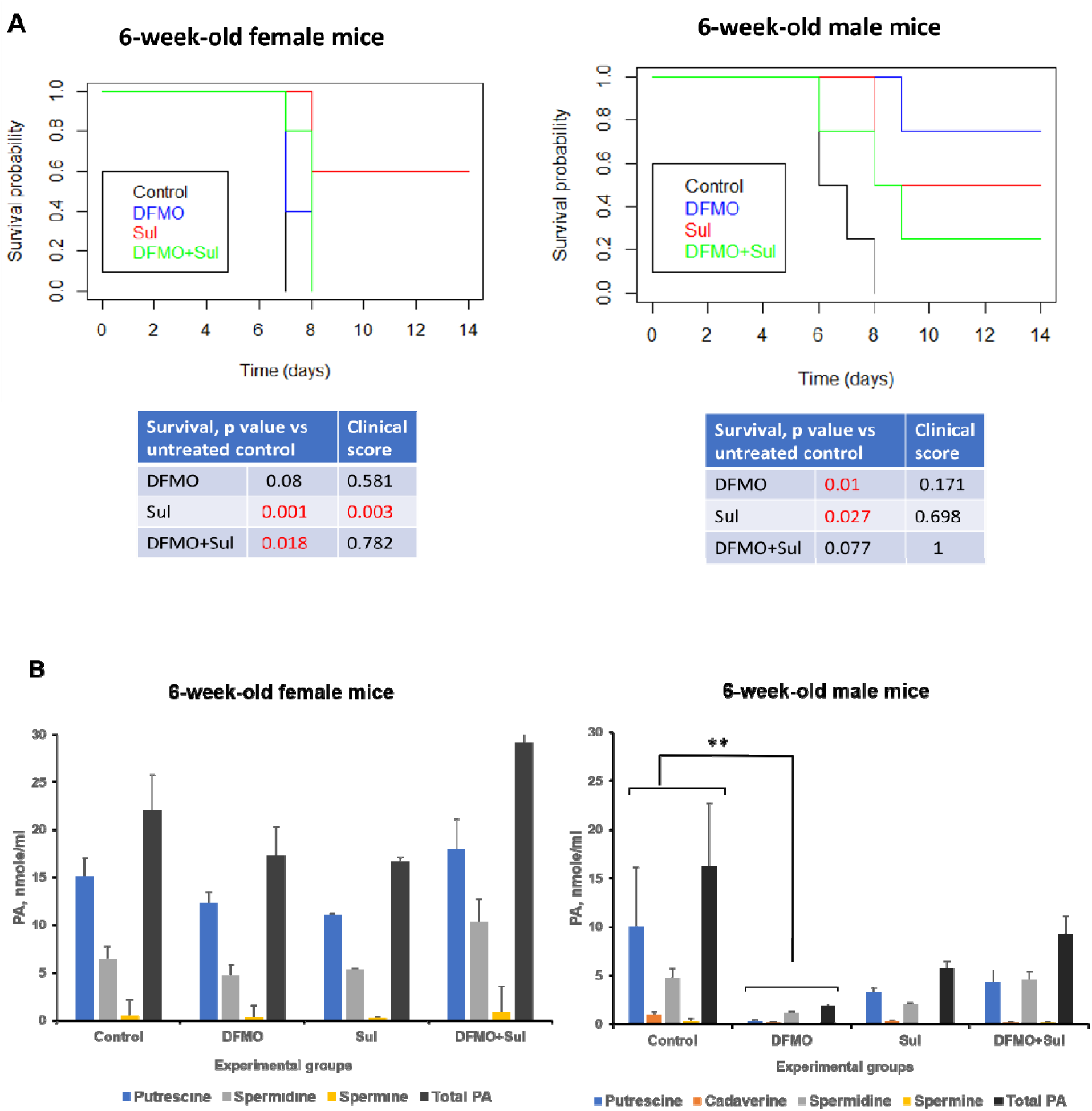
Effect of DFMO and Sulindac prophylaxis on survival and plasma polyamine content in young (6-week-old) *K18-hACE2* mice. Mice received AIN93G diet without drugs supplementation (Control) or receiving the medicated diets with 835 ppm DFMO (DFMO), 167 ppm Sulindac (Sul) or 835 ppm DFMO and 167 ppm Sulindac (DFMO+Sul) for 7 days prior to infection with 1000 PFU of SARS-CoV-2. The medicated diets were provided through the duration of the experiments (14 dpi). **A.** Kaplan-Meyer curves with p values of survival rates and clinical scores for young female and male mice. The difference in survival rate was determined using long-rank test and Cox proportion hazard test. One-way ANOVA was used to performed to compare clinical scores between treatment groups. **B**. Polyamine content in plasma of young female and male mice. The p-values for Control vs treatment groups were determined by one-way ANOVA/two-sample t-test, and a p-value p<0.05 was considered significant. **p<0.01

The Kaplan-Meyer curves analysis in young male mice showed a statistically significant increase in the survival of animals in all treatment groups compared to the survival of infected untreated group. While untreated male mice developed the morbidity symptoms by 8 dpi (mean survival 6.75 dpi), DFMO-treated group had a mean survival of 12.75 dpi (p=0.01), mice treated with Sulindac had a mean survival of 11.00 dpi (p=0.027), and DFMO/Sulindac combo group had a mean survival of 9.25 dpi, although the difference in survival did not reach statistical significance (p=0.077). The clinical scores were not significantly affected by any treatment in male mice (**Fig. 4A**, **6-week-old male mice).** The statistical analysis of young animal survival also revealed that the female mice, which were fed a Sulindac-supplemented diet, and the male mice, fed DFMO-supplemented diet, had the lowest hazard ratios compared to the control groups (0.032 and 0.044, respectively) (**Supp. Materials -Prophylaxis. Survival Analysis**).

#### Plasma polyamines analysis

Polyamine analysis in the plasma of the young female mice showed no statistically significant difference between untreated and treated groups (**Fig. 4B**, **Polyamine content**). The statistical analysis of polyamine content in the plasma samples of young male mice identified a significant reduction in putrescine (p=0.048), cadaverine (p=0.007), spermine (p=0.01), and the total polyamine content (p=0.008) in DFMO-treated mice compared to untreated mice. The Sulindac-treated male mice showed a significant reduction in the levels of cadaverin (p=0.045) and spermine (p=0.038). The males, treated with DFMO/Sulindac combination, had a significantly lower level of cadaverin (p=0.008). Still, the levels of other polyamines did not change significantly from the polyamine levels in untreated mice. The statistically significant sex-treatment interaction in the level of plasma spermine was found in 6-week-old male mice (p=0.008). The statistical analysis of polyamine content in young animals is presented in **Supplementary Materials, Prophylaxis. Polyamine content**).

##### Prophylaxis regime in aged mice

Similarly to the young mice above, 58-week-old K18-hACE2 mice were provided medicated diets for 7 days before the intranasal infection with 1000 PFU SARS-CoV-2, with continuing drug supplementation in the medicated diets for the duration of the experiment (14 dpi).

#### Survival analysis

Untreated aged female mice lived slightly longer than untreated aged male mice (mean survival time for female mice was 10.5 dpi vs 8.25 dpi for male mice). The survival rates for the untreated female mice were 0.25, while the survival rates of female mice in all treatment groups were 0.50. The DFMO/Sul combo-treated female mice had the longest mean survival time of 12 dpi. (**Fig.5A, Left panel**). The survival rate of the control untreated and DFMO-treated aged male mice was 0 at 14 days post-infection, while Sulindac-treated mice and DFMO/Sulindac combo had survival rates of 0.25 and 0.50, respectively (**Fig. 5A, right panel).** The mean survival of the infected untreated aged male mice was 8.25 dpi. The DFMO-treated male mice had a lower mean survival of 7 dpi (p=0.066). Even though the Sulindac-treated aged male mice survived longer than untreated mice (mean 10.75 dpi, p=0.097), the only group with a statistically significant increase in survival was the DFMO/Sul combo-treated group, with a mean survival of 11.75 dpi (p=0.042). The clinical scores did not significantly improve in aged male mice of any treatment group. The hazard ratio analysis of aged animals survival in all treatment groups were high, but were smaller in DFMO/Sulindac treated female and male mice, compared to the animals treated with single compounds (**Supplementary Materials. Survival Data-Prophylaxis).**

**Figure 5.**
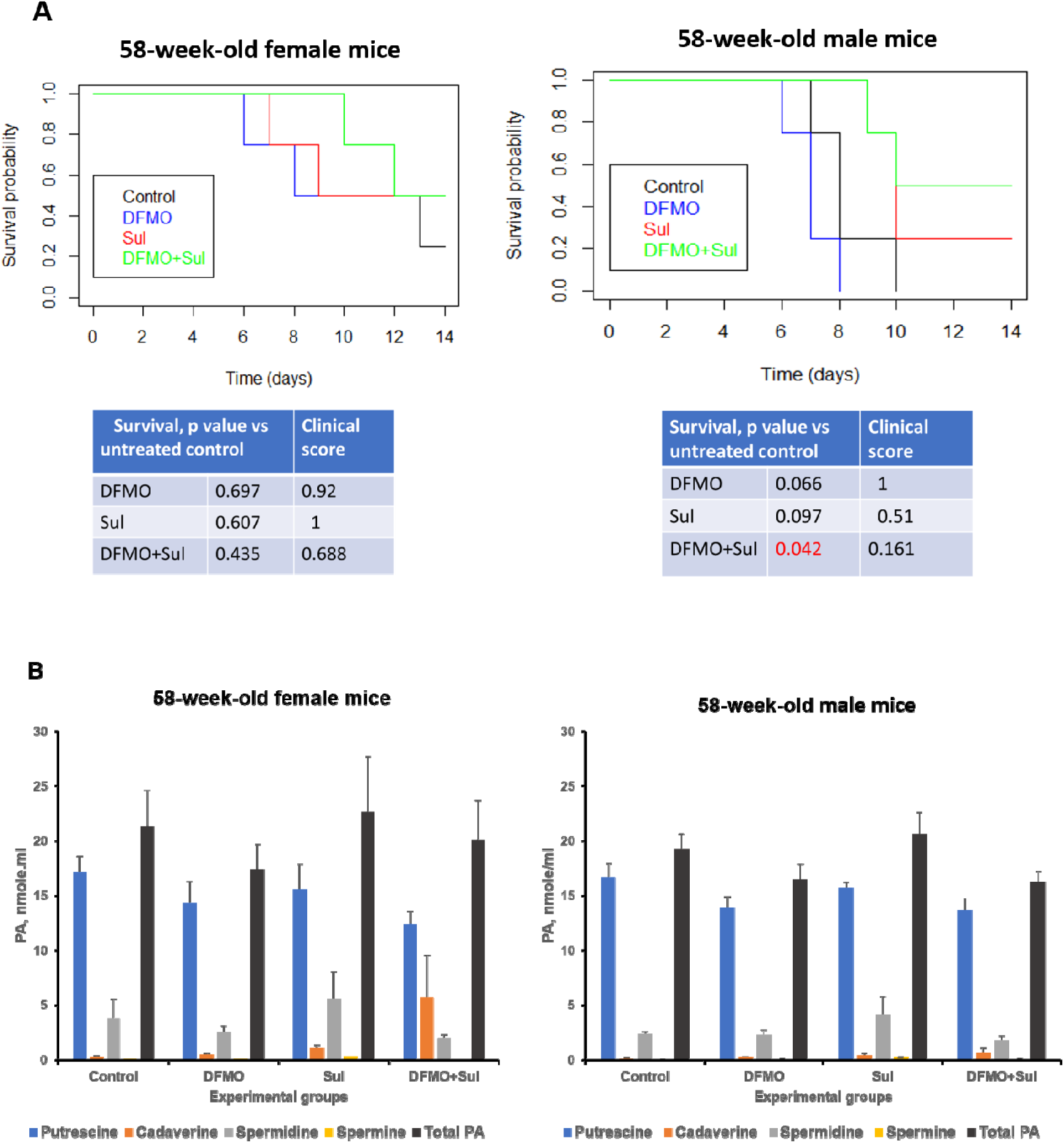
Effect of DFMO and Sulindac prophylaxis on survival and plasma polyamine content in aged (58-week-old) *K18-hACE2* mice. Mice received AIN93G diet without drugs supplementation (Control) or receiving the medicated diets with 835 ppm DFMO (DFMO), 167 ppm Sulindac (Sul) or 835 ppm DFMO and 167 ppm Sulindac (DFMO+Sul) for 7 days prior to infection with 1000 PFU of SARS-CoV-2. The medicated diets were provided throughout the duration of the experiments (14 dpi). **A.** Kaplan-Meyer curves with p values of survival rates and clinical scores for aged female and male mice. The difference in survival rates was determined using long-rank test and Cox proportion hazard test. One-way ANOVA was used to compare clinical scores between treatment groups. A p≤0.03 value between the Control and treatment groups was considered significant. **B**. Polyamine content in plasma of aged female and male mice. One-way ANOVA/two-sample t-test was performed to compare polyamine content. A p<0.05 value between Control and treatment groups was considered significant.

#### Plasma polyamine analysis

No statistically significant difference was found in the levels of plasma polyamines between the infected untreated and treated aged animals of both sexes. (**Fig. 5B** and **Supplementary Materials, Polyamine content Prophylaxis.**).

##### Treatment regime in young mice

The 6-week-old, *K18-hACE2* mice (5 per group) were placed on the AIN-93G diet for 1 week before they were infected intranasally with 1000 PFU SARS-CoV-2 and were maintained on the AIN-93G diet throughout the experiment. Treatments with DFMO, Sulindac, and DFMO/Sulindac combo in young mice were administered once daily by intragastric gavage (IG) and continued to a maximum of 10 dpi (**Fig. 3B, left panel**). The selected treatment doses of 154.17 mg/kg (mg of compound per kg of body weight) DFMO and 30.8 mg/kg of Sulindac were equivalent to the human doses tested in patients with prior colon polyps (ClinicalTrials.gov Identifier: NCT00118365) (50). Control groups in these experiments were infected untreated mice, which received the drug diluents (water-control group) and 0.5M sodium bicarbonate (Sodium group). The treatments started 24 hours post-infection, and animals were sacrificed when they met the criteria for moribundity.

#### Survival analysis

The young female mice treated with DFMO or Sulindac improved their survival rates compared to untreated ones from 0 to 0.2, and Sodium-treated female mice had a 0.6 survival rate. Female mice that received the DFMO/Sulindac combo did not show improvement in survival rate compared to the untreated (Control) group (**Fig. 6A, 6-week-old female mice**). Treatment of the young male mice with Sulindac alone or Sulindac in combination with DFMO did not improve their survival compared to the survival of the untreated (Control) mice. However, mice that received Sodium diluent or DFMO alone had a slightly improved median survival rate of 0.4 (mean survival 8 and 7 dpi, respectively) **(Fig.6A, 6-week-old male mice and Supplementary Materials. Treatment. Survival analysis).**

**Figure. 6.**
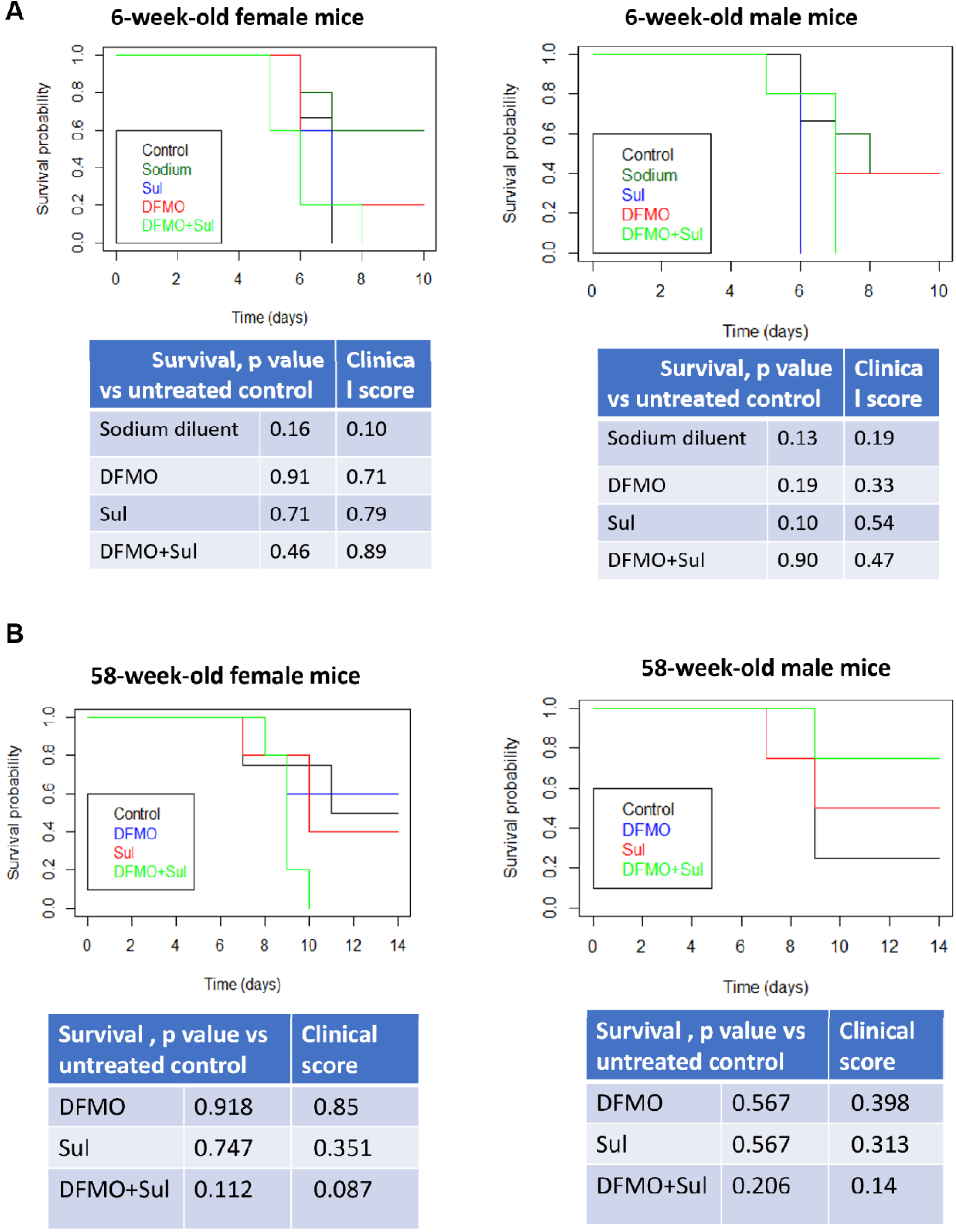
Effect of DFMO and Sulindac treatment on survival rate in young (6-week-old) and aged (58-week-old) *K18-hACE2* mice. The treatments started 24 h after animals were infected with 1000 PFU of SARS-CoV-2. **A**. Kaplan Meyer curves in young female and male mice, which received daily intragastric gavage (IG) of water-diluent (Control), 0.5M sodium bicarbonate diluent (Sodium), 31 mg/kg of Sulindac dissolved in 0.5M sodium bicarbonate (Sul), 154 mg.kg DFMO, dissolved in water (DFMO), or 31mpk Sulindac and 154 mg/kg DFMO, dissolved in 0.5M sodium bicarbonate (DFMO+Sul). The drugs were administered via daily intragastric gavage through the duration of the experiments (10 dpi). **B.** Kaplan-Meyer curves in aged female and male mice that received AIN93G diet without drugs supplementation (Control) or the medicated diets with 835 ppm DFMO (DFMO), 167 ppm Sulindac (Sul) or 835 ppm DFMO and 167 ppm Sulindac (DFMO+Sul). The medicated diets were provided through the duration of the experiments (14 dpi). The p values for survival rates and clinical scores between Control and treatment groups are presented for young and aged female and male mice. The difference in survival rates was determined using long-rank test and Cox proportion hazard test. One way ANOVA was used to compare clinical scores between treatment groups. A p<0.05 value between Control and treatment groups was considered significant.

#### Plasma Polyamine analysis

The plasma polyamine levels were not significantly altered in young mice of both sexes by the treatments compared to control mice that received sodium bicarbonate diluent, except for a significantly higher spermidine level in DFMO/Sulindac-treated young male mice (p=0.004) (**Supplementary Materials, Treatment, Polyamine content**). The infected mock-treated 6-week-old male mice (administered water by IG) had a significantly higher content of putrescine and total polyamines (p<0.0001) compared to all other experimental groups (diluent, DFMO, Sulindac, and DFMO/Sul combo). The plasma polyamine content in 6-week-old female mice was not significantly altered between the infected mock-treated group and the diluent-treated group (**Supplementary Materials, treatment**. Polyamine content).

##### Treatment regime in aged mice

The aged mice were maintained on AIN-93G diet for 7 days before the intranasal infection with 1000 PFU of SARS-CoV-2. Twenty-four hours post-infection the treatment groups were switched to the medicated diets, AIN-93G DFMO 835 ppm, AIN93G Sulindac 167 ppm, and AIN-93G DFMO/Sulindac combo. The medicated diets were provided for the duration of the experiment (14 dpi) (**Fig. 3B, right panel**).

While the untreated aged female mice had a 0.5 survival rate by day 14 post-infection, and the DFMO-treated and Sulindac-treated mice had survival rates of 0.6 and 0.4, respectively, none of the mice receiving the DFMO/Sulindac combo diet survived up to 14 dpi. The clinical score did not significantly improve by the treatments, except for the DFMO/Sulindac combo-treated animals which showed some improvement in clinical score, although the p-value did not reach statistical significance (p=0.087) (**Fig. 6B, 58-week-old female mice**). The aged male mice showed a trend of increased survival by treatments compared to the untreated group (survival rate of 0.25) with survival rates of 0.5 in DFMO- and Sulindac-treated groups and 0.75 in DFMO/Sulindac combo group (**Fig. 6B, 58-week-old male mice**). Similarly to the aged female mice, the treatments did not improve clinical scores in aged male mice. The hazard ratio of aged female and male mice treated with the DFMO-supplemented diet, had the lowest hazard ratio vs untreated (control) mice (0.902 and 0.593, respectively) (**Supplementary Materials. Survival analysis. Treatment**). The average daily consumption of food (AIN93G diet control and AIN93G-medicated diets) gradually decreased in the infected animals of all experimental groups from 3.5 g /day (9.6% of body weight consumed) on the 1^st^ dpi to 1.08 g (3.17% of body weight consumed) on 6 dpi, and this decrease correlated with the increase in the clinical score of the infected animals. This decreased food consumption in the infected animals may explain the lack of improvement in infected mice receiving medicated diets.

### Plasma polyamine analysis

The plasma polyamine content was not significantly altered in 58-week-old infected female mice receiving the medicated diets as compared to mice on the control AIN-93G diet. The 58-week-old male mice treated with Sulindac had higher plasma putrescine (p=0.02) and total polyamine levels (p<0.01) compared to the untreated animals (**Supplementary Materials).**

## Analysis of sulindac metabolites in young and aged animals on prophylaxis and treatment regimes

The NSAID sulindac is a prodrug, which is metabolized in the liver into two forms: sulindac sulfide, the non-selective inhibitor of COX-2, and sulindac sulfone, which does not have anti-inflammatory activity, but induces the polyamine catabolic enzyme SAT1 (29).

### Prophylaxis regime

Analysis of Sulindac and its metabolites in the plasma of young female and male mice treated with Sulindac alone or the DFMO/Sulindac combo is presented in **Figure 7A** and in **Supplementary Materials, Prophylaxis, Sulindac metabolites analysis)**. The young mice of both sexes had a similar level of sulindac sulfone metabolite in plasma, which was 5-fold higher than the Sulindac sulfone level in the plasma of aged mice (**Fig. 7A, left panel**). Within the category of young animals, the Sulindac-treated mice had a higher level of Sulindac in the plasma (p=0.068) compared to the DFMO/Sulindac-treated groups, although the differences did not reach statistical significance. The 6-week-old male mice treated with DFMO/Sulindac combo had on average twice higher content of Sulindac and 3 times higher content of Sulindac sulfide in the plasma, compared to the young female mice in the same group. The content of Sulindac sulfone was similar in Sulindac-treated and DFMO/Sulindac combo groups for both sexes. This observation indicates that DFMO/Sulindac combination prophylaxis in young mice resulted in the high rate of Sulindac oxidation into Sulindac Sulfone in young male mice, but not in young female mice. Plasma Sulindac and Sulindac sulfide levels were similar in the aged female and male mice treated with Sulindac or DFMO/Sulindac combo (**Fig. 7A, right panel**). However, the aged female mice treated with DFMO/Sulindac combo had a lower plasma level of Sulindac sulfone than Sulindac-treated female mice (p=0.067). Levels of Sulindac and its metabolites were similar in the aged male mice treated with Sulindac or DFMO/Sulindac combo. We also noted that aged mice of both sexes had significantly lower levels of Sulindac Sulfone metabolite compared to the young mice when treated with Sulindac alone (p<0.01 or with DFMO/Sulindac combo (p<0.03). These data indicate the possible sex and age-specific activity of Sulindac metabolizing enzymes, Sulindac reductase and microsomal cytochrome P450 system (51), in the liver of infected animals.

**Figure 7.**
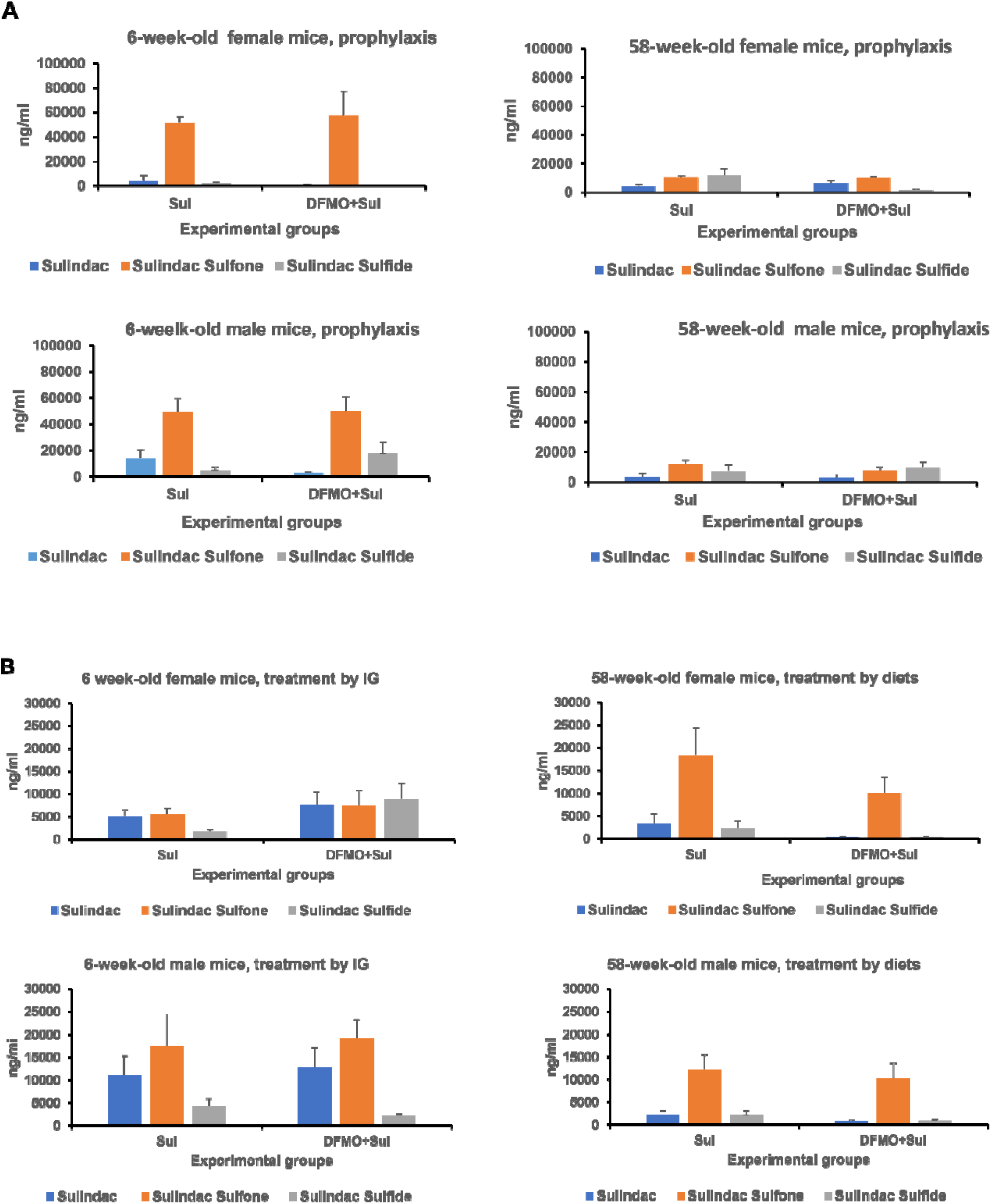
The levels of Sulindac and its metabolites Sulindac Sulfone and Sulindac Sulfide in young (6-week-old) and aged (58-week-old) *K18-hACE2* mice receiving prophylaxis or treatment regimes with Sulindac alone (Sul) or DFMO /Sul combo (DFMO+Sul). **A**. The levels of Sulindac and its metabolites Sulindac Sulfone and Sulindac Sulfide in young (6-week-old animals) (**Left panels**) and aged (58-week-old animals (**Right panels**) receiving prophylaxis regime Data was analyzed using one-way ANOVA/two-sample t-test. No statistically significant difference was found between Sulindac group and DFMO/Sul combo group within each sex. **B.** Effect of DFMO and Sulindac treatment regime on the levels of Sulindac and its metabolites Sulindac Sulfone and Sulindac Sulfide in 6-week-old animals when drugs were administered via IG (**Left panels**) and 58-week-old animals treated via medicated diets (**Right panels**) supplemented with Sulindac alone (Sul) or DFMO /Sul combo (DFMO+Sul). Data was analyzed using one-way ANOVA/two-sample t-test. No statistically significant difference was found between the Sulindac group and the DFMO/Sul combo group within each sex.

### Treatment regime

Analysis of Sulindac metabolites in young mice infected and treated with Sulindac or DFMO/Sulindac combo showed no difference in metabolite levels between the treatment groups of both sexes. The aged female mice had higher values of Sulindac sulfide and Sulindac sulfone in the Sulindac-treated group than in the DFMO/Sulindac group, with Sulindac sulfone especially elevated in the Sulindac group (p=0.09). In aged animals, the plasma level of Sulindac sulfone in the DFMO/Sulindac combo group was higher in male mice than female mice, and Sulindac sulfide was lower in male mice, compared to female mice, although their values did not reach statistical significance (p=0.078 and 0.099, respectively) (**Fig. 7B and Supplementary Materials, Treatment, Sulindac metabolites analysis**).

## Effects of DFMO and Sulindac prophylaxis and treatment on SARS-CoV-2 infectivity and viral load

To evaluate the ability of DFMO and Sulindac to suppress SARS-CoV-2 infectivity we performed a plaque-forming assay with lung tissue samples from infected mice untreated or treated with DFMO, Sulindac, and DFMO/Sulindac combo. Plaque forming assay is a quantitative method for measuring the infectious SARS-CoV-2 virus and is considered a gold standard assay for measuring virions capable of replication (34,52). Plaque-forming assay was done in cultured Vero cells infected with serial dilutions of the homogenized lung tissue samples from mice which received DFMO and Sulindac under the prophylaxis and treatment regimes. Plaques were counted after 72 hours incubation in a methylcellulose-containing (1% w/v) cell culture medium. The effect of drug treatments on SARS-CoV-2 viral load was done by measuring the N1 transcript copy number in the homogenized lung tissue of mice that received DFMO and Sulindac under prophylaxis and treatment regimes.

### Prophylaxis

We observed high variability in the results of plaque assay and N1 qPCR from the lung tissue among the infected untreated and treated mice, which were placed on the prophylaxis regime. Nevertheless, we noted the sex and age differences in the infectivity and viral load of the virus among the experimental groups. Particularly, the plaque count in the control and DFMO-treated groups was higher in the young female mice and aged male mice, and the Sulindac-alone and DFMO/Sulindac groups developed less plaques, compared to the untreated groups (**Fig. 8A**). The analysis of N1 transcript level in the lung tissue of mice on prophylaxis regime showed more than 25-fold decrease in young female mice treated with DFMO alone, while Sulindac alone and DFMO/Sulindac combo treatments did not have any significant effect on N1 transcript level in these mice. The level of N1 transcript in young male mice in all experimental groups was low and no statistically significant difference was found between control and treatment groups (**Fig.8B, 6-week-old mice**). On a contrary, a difference in the N1 transcript level was noted between aged male and female mice, with the 10 times higher N1 mRNA level in male vs female mice (**Fig.8B, 58-week-old mice**). The aged male mice treated with DFMO/Sul combination had a more than 30-fold decrease in N1 transcript level when compared to infected control male mice.

**Figure 8.**
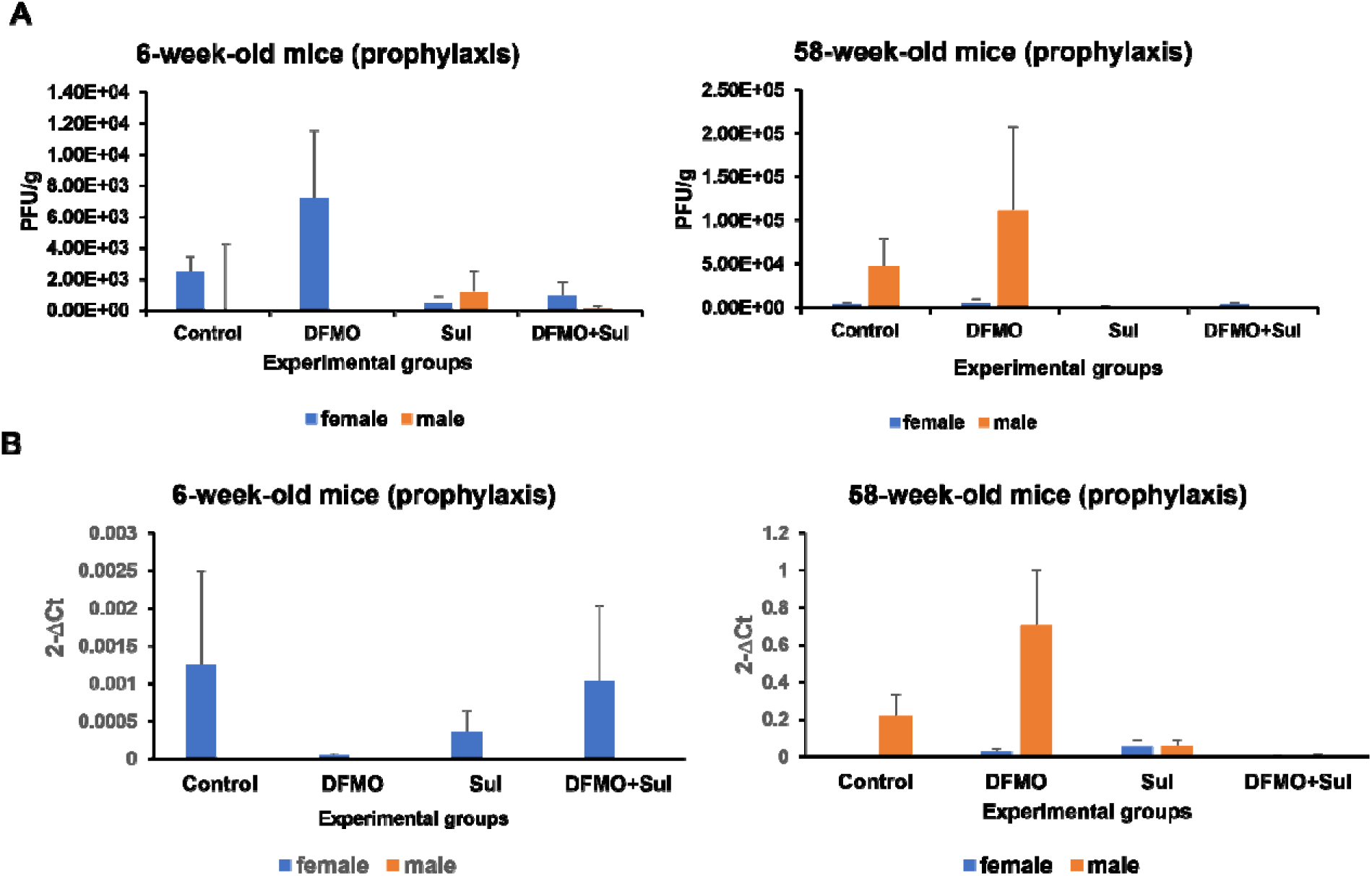
Effect of DFMO and Sulindac prophylaxis on SARS-CoV-2 infectivity and viral load in the lung tissue of infected *K18-hACE2* mice. Mice received an AIN93G diet without drugs supplementation (Control) or AIN-93G diet supplemented with 835 ppm DFMO (DFMO), 167 ppm Sulindac (Sul) or 835 ppm DFMO and 167 ppm Sulindac (DFMO+Sul) for 7 days prior to infection with 1000 PFU of SARS-CoV-2. The medicated diets were provided throughout the duration of the experiments (14 dpi). **A**. Plaque forming assay results in the lung tissue of young (6-week-old mice) and aged (58-week-old) female and male mice. The results are shown as a number of plaque-forming units per g of lung tissue (PFU/g). **B**. Analysis of SARS-CoV-2 N1 gene expression in the lung tissue of infected control and treated experimental groups. Data is presented as the normalized average N1 gene expression per group. Data was analyzed using ANOVA single-factor test. No statistically significant difference was found between the control and treatment groups. C.

### Treatment

The DFMO, Sulindac and DFMO/Sul combo treatments by IG resulted in the lower infectivity of the lung tissue in young male mice. The results of the plaque-forming assays performed on the lung tissue of young and aged mice were inconclusive because the lung tissue of infected untreated control female and male mice generated a lower number of plaque-forming units than the lung tissue of treated animals (**Fig. S4**). Overall, these data suggest that the prophylaxis regime of DFMO and Sulindac decreased the infectious SARS-CoV-2 virus level in the primary target tissue (lung), with a level of protection against lethal SARS-CoV-2 infection.

## Discussion

Viruses rely on host metabolism to propagate, therefore it is important to identify anti-viral drugs that would interfere with host cellular functions that are essential to the viral life cycle. There is solid experimental evidence that inhibition of polyamine biosynthesis can suppress the replication of different viruses (9). The novel SARS-CoV-2 virus has unique genome differences compared to other coronaviruses, therefore it was important to evaluate the antiviral activity of polyamine inhibition against SARS-CoV-2.

DFMO is an inhibitor of the ODC1 enzyme, which converts the amino acid ornithine into diamine putrescine, a precursor of polyamines spermidine and spermine. As an inhibitor of polyamine synthesis, DFMO has been widely used not only for the characterization of polyamine function in mammalian cells but as a clinical drug. DFMO is approved by the US Food and Drug Administration for the treatment of trypanosomiasis (53,54), hirsutism (55) and, recently, for treating neuroblastoma patients (www.IWILFIN.com). Sulindac is an FDA-approved NSAID. Its active metabolites, sulindac sulfide, and sulindac sulfone, have anti-inflammatory and polyamine catabolism-inducing activities, respectively. The DFMO/Sulindac drug combination was successfully tested for its antitumor activity in the animal model of Familial Adenomatosis Polyposis (30). The human clinical trial demonstrated that DFMO and Sulindac as a combination treatment is safe and remarkably effective in preventing the occurrence of colorectal adenomas in patients with prior colon polyps (50). The drug combination can be safely administered as oral medications in an outpatient setting at doses that reduce polyamine levels in adults as young as 18 years of age.

The rationale for the current research was to determine the susceptibility of the SARS-CoV-2 virus to DFMO and Sulindac treatment for potential use as an alternative approach to combat COVID-19. To address the efficacy of DFMO and Sulindac as antiviral agents, we initially performed *in vitro* analysis of the DFMO/Sulindac combo in the infected Vero, Calu-3, and Caco-2 cell lines. We determined that the DFMO/Sulindac drug combination acted synergistically to suppress SARS-CoV-2 viral load when cells were pretreated with the drugs for 48 hours before exposure to the virus. The additive effect of the drug combination was observed when the treatment started after cells were infected. This finding indicates that polyamine depletion in cells before infection improves the DFMO and Sulindac combined antiviral effect. The expression of SARS-CoV-2 Spike and Nucleocapsid (N1) were significantly suppressed in the Calu-3 cell line treated with Sulindac alone or DFMO/Sulindac combo (p<0.0001). SARS-CoV-2-infected Calu-3 cells exhibited a significant induction of the mRNA levels of key polyamine metabolic enzymes *ODC1*, *SAT1,* and *SMOX.* DFMO and Sulindac treatments as single agents, or in combination, suppressed *SAT1* and *SMOX* gene expression in infected cells. The tested compounds did not alter the ODC mRNA level because DFMO inhibits ODC enzymatic activity, not ODC transcription (19).

Next, we tested the DFMO/Sulindac prophylaxis and therapy regimes against SARS-CoV-2 infection *in vivo* using the *C57BL/6J K18-hACE2* transgenic mouse model. *K18-hACE2* mice are highly susceptible to SARS-CoV-2 infection and upon intranasal infection with the SARS-CoV-2 virus faithfully recapitulate the signs of a severe COVID-19 in humans, including body weight loss, rapid breathing, fever, and inactivity. The infected mice on average reach criteria for euthanasia within 5-8 days post-infection when infected with SARS-CoV-2 at 1×10^5^ PFU (56). In this study, *K18-hACE2* mice were infected with 1000 PFU and were monitored for 14 days. The analysis of plasma polyamine levels in infected untreated *K18-hACE2* mice revealed a sex-specific difference, with more significantly altered polyamine levels in male mice as compared to female mice. Particularly, uninfected young male mice had a lower polyamine content than female mice, whereas SARS-CoV-2 virus significantly induced polyamine content, specifically putrescine, in infected male mice. The uninfected aged male and female mice had comparable plasma polyamine levels, but SARS-CoV-2 infection suppressed polyamine content in the infected aged male mice. The observed sex-specific differences in polyamine levels in response to SARS-CoV-2 infection are important for determining the antiviral efficacy of DFMO and Sulindac. The survival rates in animals on the prophylaxis regime demonstrated striking sex- and age-specific outcomes. The prophylaxis with DFMO and Sulindac as the single agents was more effective in young male mice by increasing survival rates at 14 days post-infection (dpi) from 0 to 0.75 (DFMO), and from 0 to 0.5 (Sulindac). The DFMO/Sul combo was marginally effective in young male mice by increasing their survival rate by 0.25 (p=0.077). At the same time, the DFMO/Sul combo diet significantly improved the survival of aged male mice (p=0.042), although the individual drugs did not. The young female mice benefited from the prophylaxis regime with Sulindac alone (increase in survival rate from 0 to 0.6, p=0.001, and a statistically significant increase in clinical score, p=0.003). The young female mice also showed a significant increase in survival rates on the DFMO/Sul combo diet (p=0.018). Low hazard ratios were observed in young DFMO-fed male mice and young Sulindac-fed female mice, which suggests a shortened duration of illness in these experimental groups. The survival rates of aged female mice did not improve significantly by any prophylaxis treatments (DFMO, Sulindac, or DFMO/Sul combo) because the infected untreated mice had a high mean survival time of 10.5 days. The survival rate analysis in animals on the treatment regime showed similar median survival rates in young male and female mice treated with DFMO, Sulindac, or their combination. The treatment regimes slightly improved the median survival times of the aged male mice, with DFMO/Sul combination treatment being the most effective in aged male mice (survival rate increased from 0.25 to 0.75), while aged female mice did not significantly improve on any treatments.

Analysis of plasma polyamine levels in animals from different treatment groups showed that DFMO-treated young male mice had 8 times less polyamine concentration compared to all other groups. A statistically significant sex-treatment interaction was found in the plasma spermine level in 6-week-old male mice. Plasma polyamine levels were not significantly altered by any treatment in aged mice. The analysis of sulindac metabolites revealed a 4-fold higher concentration of Sulindac sulfide in the plasma of male mice as compared to female mice. Sulindac oxidation to sulindac sulfone is catalyzed primarily by the microsomal cytochrome P450 (P450) system, and sulindac sulfide is generated via reduction by the thioredoxin system. The observed differences in metabolite levels can be due to the different activities of these enzyme complexes in male and female mice, which can change with age. Sulindac sulfide inhibits COX-1 and COX-2 >48-fold more potently than sulindac sulfone metabolite, resulting in the suppression of prostaglandin (PG) levels (57), but also can cause hepatic toxicity (58). Sulindac has been shown to inhibit the activities of the two isoforms (COX-1 and COX-2) of cyclooxygenase enzyme, and to induce apoptosis through both COX-dependent (59) and COX-independent mechanisms (29).

Both innate and adaptive immunity, which is regulated by sex hormones and is affected by aging, may play a role in the sex-and age-dependent differences observed in our study (60,61). Additionally, polyamines can modulate the expression of pro-inflammatory genes and cytokine synthesis (62) and can induce an autoimmune response (63). The sex difference in plasma polyamine levels between infected control and treated young animals requires further investigation of the immune response against COVID-19 elicited by the DFMO prophylaxis regime.

Overall, preclinical *in vivo* studies demonstrated a selective, protective, age- and sex-dependent antiviral effect of DFMO and Sulindac treatment in young and aged animals. We found that plasma polyamine levels were reduced in male mice on a prophylaxis regime, which could be associated with the observed viral load suppression and reduction of inflammation.

The reduced viral loads were associated with lessening the severity of clinical symptoms, pathology, and lethality in SARS-CoV-2-infected *K18 hACE2* mice. Interestingly, SARS-CoV-2-infected *K18 hACE2* mice exhibited both age- and sex-dependent differences in animals treated with the DFMO/Sulindac combo, but only sex-dependent differences were found in Sulindac metabolism.

This study also demonstrated some limitations of the *K18hACE2* transgenic mouse model. We ensured the reproducibility of animal studies by using a statistically justified number of animals representing both sexes per treatment group, per time point, and three technical variables for each analysis to account for tissue heterogeneity. Nevertheless, we observed variability in viral loads among the infected mice and less than desired suppression of viral infection by DFMO/Sul in some experiments. The observed variability could be due to variability in the activity of the K18-promoter, which controls the expression of hACE2 in target organs (lung, brain, intestine, heart, liver kidney, spleen) of *K18 hACE2* transgenic animals. Additionally, the development of anosmia in the infected *K18hACE2* (56) could suppress their attraction to food, which may also have contributed to the variability of the reported data.

Nevertheless, our studies showed the promising effect of DFMO and Sulindac prophylaxis of viral infection with a specific protective age- and sex-dependent antiviral effect of DFMO as a single agent in young animals and DFMO/Sulindac combination in aged animals. The results of our animal studies recapitulated the epidemiological reports regarding the higher vulnerability of men as compared to women to COVID-19-related mortality. They also demonstrated that the administration of DFMO and Sulindac could be useful antiviral agents for personalized protection and treatment of SARS-CoV-2 infection. Our studies provide sufficient pre-clinical efficacy data to pursue additional evaluation of DFMO and Sulindac as a viable preventive approach against SARS-CoV-2 infection and for the treatment of “long COVID” symptoms.

## Acknowledgements.

The presented work was supported by the National Center for Advancing Translational Sciences under award numbers UG3TR003597 (to N.A. Ignatenko and D.G. Besselsen) and UH3TR003597 (to N.A. Ignatenko and C. Bime).

## Abbreviations and acronyms

COVID-19: COronaVIrus Disease 2019
SARS-CoV-2: Severe Acute Respiratory Syndrome CoronaVirus 2
GEM: genetically engineered mouse
NSAID: non-steroidal anti-inflammatory drug
DFMO: α-difluoromethylornithine
Sul: Sulindac
COX2: cyclooxygenase 2
ODC1: ornithine decarboxylase
SAT1: spermidine/spermine N1 acetyltransferase
SMOX: spermine oxidase
N1: Nucleocapsid gene
DMEM: Dulbecco’s modified Eagle medium
MOI: multiplicity of infection
PFU: plaque-forming units
HPLC: high-performance liquid chromatography.

