## Supplemental Materials for "Preclinical evaluation of the efficacy of α−Difluoromethylornithine and Sulindac against SARS-CoV-2 infection"

SUPPLEMENTARY MATERIALS

**Figures.**

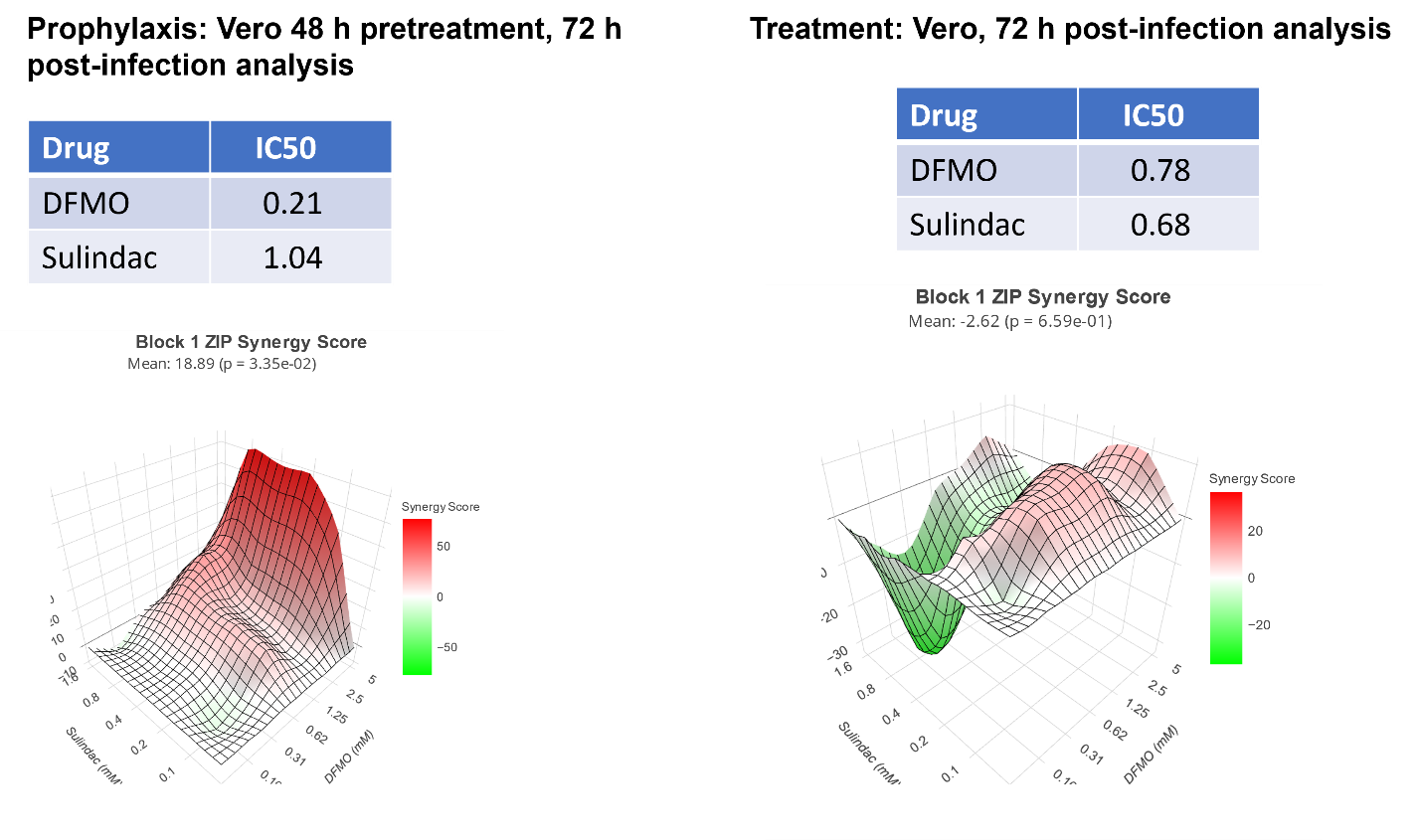

**Figure S1.** Results of Synergy Finder analysis of DFMO/Sul in Vero cell line, infected with MOI 0.05. Cells were incubated with different concentrations of DFMO and Sulindac either for 48 h (Prophylaxis) before infection or drugs were added 24 h after the virus was added (treatment) as described in detail in the Materials and Methods section. In both conditions N1 transcript level was measured by the qPCR 72h post-infection.

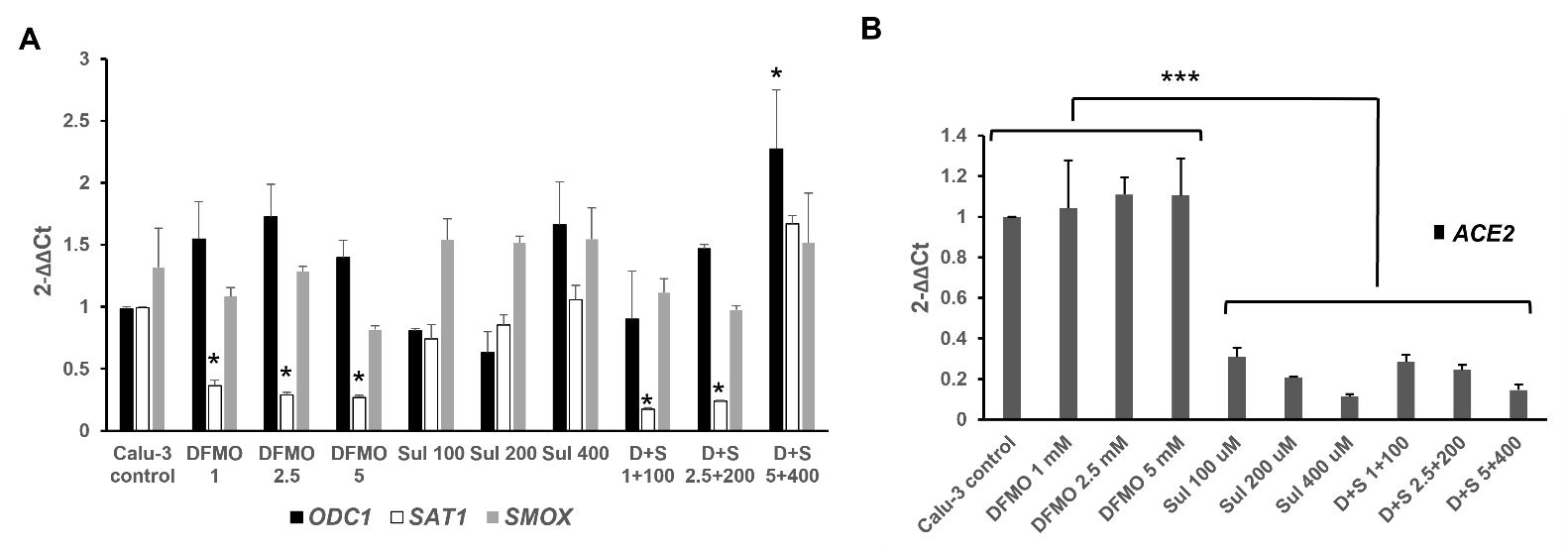

**Figure S2**. Gene expression analysis in uninfected Calu-3 lung adenocarcinoma cells treated with DFMO and Sulindac as single agents and in combinations by qPCR. **A.** Fold change in polyamine metabolic genes expression in Calu-3 cells incubated with the various concentrations of DFMO (1 mM, 2.5 mM, 5 mM), Sulindac (Sul) (100 μM , 200 μM, 400 μM), and DFMO/Sulindac combo (D+S) at 72 h .

**B.** Fold change in expression of *ACE2* mRNA in Calu-3 cells incubated with the various concentrations of DFMO (1 mM, 2.5 mM, 5 mM), Sulindac (Sul) (100 μM, 200 μM, 400 μM), and DFMO/Sulindac combo (D+S) for 72 h.

Data was analyzed using ANOVA-single factor test. *p<0.02, ***p <0.001. Results representative of three independent experiments are shown.

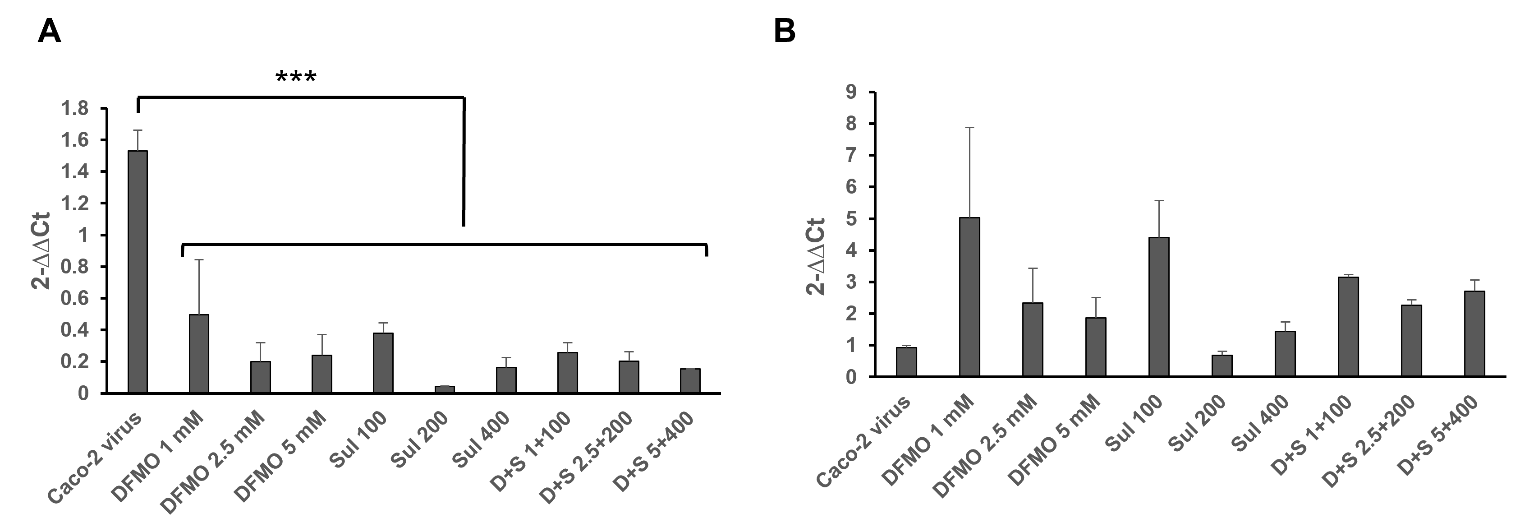

**Figure S3**. Analysis of viral gene expression in the Caco-2 colon adenocarcinoma cell line infected with SARS-CoV-2 virus at MOI 0.05 (Caco-2 virus) and treated with various concentrations of DFMO and Sulindac for 72h. A. Fold change in N1 mRNA. B. Fold change in Spike mRNA level. Data was analyzed using ANOVA-single factor test. ***p<0.001.

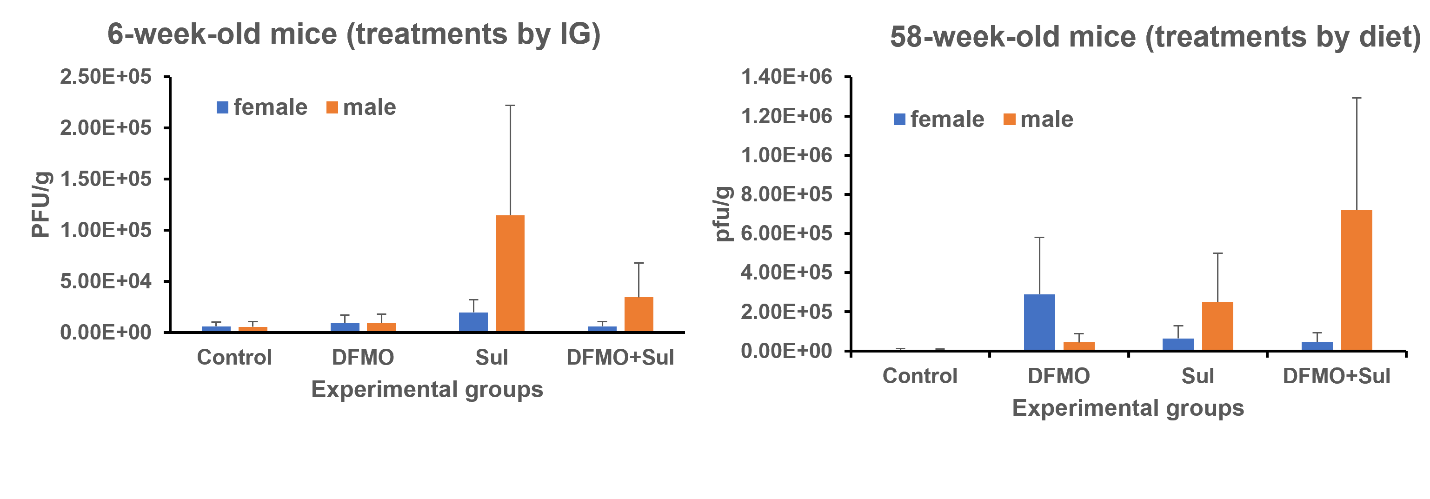

**Figure S4.** Effect of DFMO and Sulindac treatment regime on the plaque-forming ability in lung tissue of the infected *K18-hACE2* mice. Plaque-forming assay results in the lung tissue of young (6-week-old mice) and aged (58-week-old) female and male mice. The results are shown as a number of plaque-forming units per g of lung tissue (PFU/g).

**Tables.**

**PROPHYLAXIS REGIME. Survival Analysis in young and aged mice**

### Table 1: Summary of survival data for 6 week-old female mice (control vs medicated diets)

| **Group** | **N** | **Survival^a^** | **Median (days)^b^** | **Mean (days)^c^** | **HR (vs. Control)^d^** | **2.5%** | **97.5%** | **p-value** |
| --- | --- | --- | --- | --- | --- | --- | --- | --- |
| Control | 5 | 0.0 | 7 | 7.0 |  |  |  |  |
| DFMO | 5 | 0.0 | 7 | 7.4 | 0.27 | 0.062 | 1.172 | 0.08 |
| Sul | 5 | 0.6 | NA | 11.6 | 0.03 | 0.004 | 0.23 | 0.001 |
| DFMO+Sul | 5 | 0.0 | 8 | 7.8 | 0.16 | 0.033 | 0.726 | 0.018 |

### ^a^survival rate at 14 days

^b^median survival time derived from Kaplan-Meier curves

^c^mean survival time derived from Kaplan-Meier curves

^d^hazard ratio derived from Cox regression

### Table 2: Summary of clinical score for 6 week-old female mice (control vs medicated diets)

| **Group** | **N** | **Mean** | **SD** | **p-value (vs. Control)^a^** |
| --- | --- | --- | --- | --- |
| Control | 5 | 8.2 | 1.30 |  |
| DFMO | 5 | 9.0 | 1.41 | 0.581 |
| Sul | 5 | 3.2 | 3.96 | 0.003 |
| DFMO+Sul | 5 | 8.6 | 0.89 | 0.782 |

### ^a^derived from two-sample t test

**Table 3: Summary of survival data for 6 week-old male mice (control vs medicated diets)**

| **Group** | **N** | **Survival^a^** | **Median (days)^b^** | **Mean (days)^c^** | **HR (vs. Control)^d^** | **2.5%** | **97.5%** | **p-value** |
| --- | --- | --- | --- | --- | --- | --- | --- | --- |
| Control | 4 | 0.00 | 6.5 | 6.75 |  |  |  |  |
| DFMO | 4 | 0.75 | NA | 12.75 | 0.04 | 0.004 | 0.476 | 0.01 |
| Sul | 4 | 0.50 | 8.0 | 11.00 | 0.12 | 0.019 | 0.789 | 0.027 |
| DFMO+Sul | 4 | 0.25 | 8.5 | 9.25 | 0.22 | 0.041 | 1.175 | 0.077 |

### ^a^survival rate at 14 days

^b^median survival time derived from Kaplan-Meier curves

^c^mean survival time derived from Kaplan-Meier curves

### ^d^hazard ratio derived from Cox regression

### Table 4: Summary of clinical score for 6 week-old male mice (control vs medicated diets)

| **Group** | **N** | **Mean** | **SD** | **p-value (vs. Control)^a^** |
| --- | --- | --- | --- | --- |
| Control | 4 | 7.75 | 0.96 |  |
| DFMO | 4 | 5.00 | 2.71 | 0.171 |
| Sul | 4 | 7.00 | 3.16 | 0.698 |
| DFMO+Sul | 4 | 7.75 | 3.20 | 1 |

### ^a^derived from two-sample t test

### Table 5: Interaction effect between treatment and sex for 6-week-old mice

Interaction p-value

0.081

### Table 6: Summary of survival data for 58 week-old female mice(control vs medicated diets)

| **Group** | **N** | **Survival^a^** | **Median (days)^b^** | **Mean (days)^c^** | **HR (vs. Control)^d^** | **2.5%** | **97.5%** | **p-value** |
| --- | --- | --- | --- | --- | --- | --- | --- | --- |
| Control | 4 | 0.25 | 10.5 | 10.5 |  |  |  |  |
| DFMO | 4 | 0.50 | 8.0 | 10.5 | 0.70 | 0.117 | 4.199 | 0.697 |
| Sul | 4 | 0.50 | 9.0 | 11.0 | 0.63 | 0.104 | 3.748 | 0.607 |
| DFMO+Sul | 4 | 0.50 | 12.0 | 12.5 | 0.49 | 0.081 | 2.951 | 0.435 |

### ^a^survival rate at 14 days

^b^median survival time derived from Kaplan-Meier curves

^c^mean survival time derived from Kaplan-Meier curves

### ^d^hazard ratio derived from Cox regression

### Table 7: Summary of clinical score for 58 week-old female mice (control vs medicated diets)

| **Group** | **N** | **Mean** | **SD** | **p-value (vs. Control)^a^** |
| --- | --- | --- | --- | --- |
| Control | 4 | 6.25 | 3.59 |  |
| DFMO | 4 | 6.00 | 2.45 | 0.92 |
| Sul | 4 | 6.25 | 4.43 | 1 |
| DFMO+Sul | 4 | 5.25 | 2.99 | 0.688 |

### ^a^derived from two-sample t test

**Table 8: Summary of survival data for 58 week-old male mice (control vs medicated diets)**

| **Group** | **N** | **Survival^a^** | **Median (days)^b^** | **Mean (days)^c^** | **HR (vs. Control)^d^** | **2.5%** | **97.5%** | **p-value** |
| --- | --- | --- | --- | --- | --- | --- | --- | --- |
| Control | 4 | 0.00 | 8 | 8.25 |  |  |  |  |
| DFMO | 4 | 0.00 | 7 | 7.00 | 4.70 | 0.903 | 24.501 | 0.066 |
| Sul | 4 | 0.25 | 10 | 10.75 | 0.27 | 0.059 | 1.263 | 0.097 |
| DFMO+Sul | 4 | 0.50 | 10 | 11.75 | 0.16 | 0.029 | 0.937 | 0.042 |

### ^a^survival rate at 14 days

^b^median survival time derived from Kaplan-Meier curves

^c^mean survival time derived from Kaplan-Meier curves

### ^d^hazard ratio derived from Cox regression

### Table 9: Summary of clinical score for 58 week-old male mice (control vs medicated diets)

| **Group** | **N** | **Mean** | **SD** | **p-value (vs. Control)^a^** |
| --- | --- | --- | --- | --- |
| Control | 4 | 9.00 | 0.82 |  |
| DFMO | 4 | 9.00 | 0.82 | 1 |
| Sul | 4 | 7.75 | 3.40 | 0.51 |
| DFMO+Sul | 4 | 6.25 | 3.78 | 0.161 |

### ^a^derived from two-sample t test

### Table 10: Interaction effect between treatment and sex for 58 week-old mice

Interaction p-value

0.287

**PROPHYLAXIS REGIME. Plasma Polyamine Analysis in young and aged mice**

**Table 1: Summary of Putrescine by treatment and sex for 6-week-old mice**

| **Treatment** | **N(Female)** | **Mean(Female)** | **SD(Female)** | **p-value (vs. Control)** | **N(Male)** | **Mean(Male)** | **SD(Male)** | **p-value (vs. Control)** | **Interaction p-value** |
| --- | --- | --- | --- | --- | --- | --- | --- | --- | --- |
| Control | 5 | 15.08 | 4.36 |  | 3 | 3.42 | 2.20 |  | 0.525 |
| DFMO | 5 | 12.32 | 2.84 | 0.418 | 4 | 0.35 | 0.29 | 0.048 |  |
| Sul | 5 | 11.08 | 3.76 | 0.247 | 4 | 3.28 | 1.08 | 0.924 |  |
| DFMO+Sul | 5 | 17.97 | 8.34 | 0.398 | 4 | 4.34 | 2.74 | 0.518 |  |

Note: p-value (vs. Control) by Sex was derived from one-way ANOVA and the interaction p-value between Treatment and Sex was derived from two-way ANOVA with the interaction terms between Treatment and Sex indicators

**Table 2: Summary of Cadaverine by treatment for 6-week-old male mice**

| **Treatment** | **N(Male)** | **Mean(Male)** | **SD(Male)** | **p-value (vs. Control)** |
| --- | --- | --- | --- | --- |
| Control | 3 | 0.69 | 0.39 |  |
| DFMO | 4 | 0.21 | 0.09 | 0.007 |
| Sul | 4 | 0.36 | 0.05 | 0.045 |
| DFMO+Sul | 4 | 0.23 | 0.14 | 0.008 |

Note: p-value (vs. Control) by Sex was derived from one-way ANOVA.

**Table 3: Summary of Spermidine by treatment and sex for 6 week-old mice**

| **Treatment** | **N(Female)** | **Mean(Female)** | **SD(Female)** | **p-value (vs. Control)** | **N(Male)** | **Mean(Male)** | **SD(Male)** | **p-value (vs. Control)** | **Interaction p-value** |
| --- | --- | --- | --- | --- | --- | --- | --- | --- | --- |
| Control | 5 | 6.47 | 2.58 |  | 3 | 0.41 | 0.64 |  | 0.274 |
| DFMO | 5 | 4.73 | 2.53 | 0.507 | 4 | 0.04 | 0.02 | 0.112 |  |
| Sul | 5 | 5.37 | 2.71 | 0.674 | 4 | 0.04 | 0.02 | 0.114 |  |
| DFMO+Sul | 5 | 10.34 | 6.75 | 0.152 | 4 | 0.19 | 0.14 | 0.319 |  |

Note: p-value (vs. Control) by Sex was derived from one-way ANOVA and the interaction p-value between Treatment and Sex was derived from two-way ANOVA with the interaction terms between Treatment and Sex indicators

**Table 4: Summary of Spermine by treatment and sex for 6 week-old mice**

| **Treatment** | **N(Female)** | **Mean(Female)** | **SD(Female)** | **p-value (vs. Control)** | **N(Male)** | **Mean(Male)** | **SD(Male)** | **p-value (vs. Control)** | **Interaction p-value** |
| --- | --- | --- | --- | --- | --- | --- | --- | --- | --- |
| Control | 5 | 0.47 | 0.34 |  | 3 | 4.68 | 2.50 |  | 0.008 |
| DFMO | 5 | 0.29 | 0.27 | 0.564 | 4 | 1.20 | 0.34 | 0.010 |  |
| Sul | 5 | 0.24 | 0.16 | 0.461 | 4 | 2.02 | 0.43 | 0.038 |  |
| DFMO+Sul | 5 | 0.88 | 0.88 | 0.212 | 4 | 4.54 | 1.88 | 0.905 |  |

Note: p-value (vs. Control) by Sex was derived from one-way ANOVA and the interaction p-value between Treatment and Sex was derived from two-way ANOVA with the interaction terms between Treatment and Sex indicators

**Table 5: Summary of total polyamines by treatment and sex for 6-week-old mice**

| **Treatment** | **N(Female)** | **Mean(Female)** | **SD(Female)** | **p-value (vs. Control)** | **N(Male)** | **Mean(Male)** | **SD(Male)** | **p-value (vs. Control)** | **Interaction p-value** |
| --- | --- | --- | --- | --- | --- | --- | --- | --- | --- |
| Control | 5 | 22.03 | 7.04 |  | 3 | 9.19 | 4.29 |  | 0.558 |
| DFMO | 5 | 17.33 | 5.34 | 0.397 | 4 | 1.80 | 0.47 | 0.008 |  |
| Sul | 5 | 16.69 | 6.06 | 0.337 | 4 | 5.71 | 1.54 | 0.152 |  |
| DFMO+Sul | 5 | 29.19 | 13.28 | 0.203 | 4 | 9.29 | 4.17 | 0.968 |  |

Note: p-value (vs. Control) by Sex was derived from one-way ANOVA and the interaction p-value between Treatment and Sex was derived from two-way ANOVA with the interaction terms between Treatment and Sex indicators

**Table 6: Summary of Sulindac by treatment and sex for 6-week-old mice**

| **Treatment** | **N(Female)** | **Mean(Female)** | **SD(Female)** | **p-value (vs. Control)** | **N(Male)** | **Mean(Male)** | **SD(Male)** | **p-value (vs. Control)** | **Interaction p-value** |
| --- | --- | --- | --- | --- | --- | --- | --- | --- | --- |
| Sul | 5 | 4422.61 | 3834.84 |  | 4 | 14069.74 | 12920.02 |  | 0.231 |
| DFMO+Sul | 5 | 771.212 | 472.31 | 0.068 | 4 | 2860.408 | 1459.87 | 0.135 |  |

**Table 7: Summary of Sulindac Sulfone by treatment and sex for 6 week-old mice**

| **Treatment** | **N(Female)** | **Mean(Female)** | **SD(Female)** | **p-value (vs. Control)** | **N(Male)** | **Mean(Male)** | **SD(Male)** | **p-value (vs. Control)** | **Interaction p-value** |
| --- | --- | --- | --- | --- | --- | --- | --- | --- | --- |
| Sul | 5 | 51319.37 | 11459.58 |  | 4 | 49018.98 | 20427.87 |  | 0.856 |
| DFMO+Sul | 5 | 57116.29 | 45160.93 | 0.788 | 4 | 49830.46 | 21678.88 | 0.958 |  |

**Table 8: Summary of Sulindac Sulfide by treatment and sex for 6 week-old mice**

| **Treatment** | **N(Female)** | **Mean(Female)** | **SD(Female)** | **p-value (vs. Control)** | **N(Male)** | **Mean(Male)** | **SD(Male)** | **p-value (vs. Control)** | **Interaction p-value** |
| --- | --- | --- | --- | --- | --- | --- | --- | --- | --- |
| Sul | 5 | 2346.27 | 1875.39 |  | 4 | 4760.514 | 4483.8 |  | 0.106 |
| DFMO+Sul | 5 | 671.91 | 532.70 | 0.091 | 4 | 17385.09 | 18187.49 | 0.226 |  |

Note: p-value (vs. Control) by Sex was derived from one-way ANOVA and the interaction p-value between Treatment and Sex was derived from two-way ANOVA with the interaction terms between Treatment and Sex indicators

**Table 9: Summary of Putrescine by treatment and sex for 58 week-old mice**

| **Treatment** | **N(Female)** | **Mean(Female)** | **SD(Female)** | **p-value (vs. Control)** | **N(Male)** | **Mean(Male)** | **SD(Male)** | **p-value (vs. Control)** | **Interaction p-value** |
| --- | --- | --- | --- | --- | --- | --- | --- | --- | --- |
| Control | 4 | 17.16 | 2.90 |  | 4 | 16.69 | 2.51 |  | 0.804 |
| DFMO | 6 | 15.65 | 4.70 | 0.593 | 5 | 13.90 | 2.16 | 0.063 |  |
| Sul | 6 | 15.62 | 5.59 | 0.586 | 4 | 15.73 | 1.02 | 0.520 |  |
| DFMO+Sul | 6 | 12.37 | 2.91 | 0.102 | 4 | 13.68 | 2.15 | 0.058 |  |

Note: p-value (vs. Control) by Sex was derived from one-way ANOVA and the interaction p-value between Treatment and Sex was derived from two-way ANOVA with the interaction terms between Treatment and Sex indicators

**Table 10: Summary of Cadaverine by treatment for 58 week-old mice**

| **Treatment** | **N(Female)** | **Mean(Female)** | **SD(Female)** | **p-value (vs. Control)** | **N(Male)** | **Mean(Male)** | **SD(Male)** | **p-value (vs. Control)** | **Interaction p-value** |
| --- | --- | --- | --- | --- | --- | --- | --- | --- | --- |
| Control | 4 | 0.28 | 0.20 |  | 4 | 0.12 | 0.10 |  | 0.443 |
| DFMO | 6 | 0.48 | 0.33 | 0.952 | 5 | 0.22 | 0.14 | 0.772 |  |
| Sul | 6 | 1.14 | 0.48 | 0.791 | 4 | 0.45 | 0.31 | 0.361 |  |
| DFMO+Sul | 6 | 5.72 | 9.42 | 0.107 | 4 | 0.65 | 0.95 | 0.146 |  |

Note: p-value (vs. Control) by Sex was derived from one-way ANOVA and the interaction p-value between Treatment and Sex was derived from two-way ANOVA with the interaction terms between Treatment and Sex indicators

**Table 11: Summary of Spermidine by treatment and sex for 58 week-old mice**

| **Treatment** | **N(Female)** | **Mean(Female)** | **SD(Female)** | **p-value (vs. Control)** | **N(Male)** | **Mean(Male)** | **SD(Male)** | **p-value (vs. Control)** | **Interaction p-value** |
| --- | --- | --- | --- | --- | --- | --- | --- | --- | --- |
| Control | 4 | 3.77 | 3.49 |  | 4 | 2.40 | 0.32 |  | 0.946 |
| DFMO | 6 | 2.59 | 1.48 | 0.613 | 5 | 2.30 | 0.98 | 0.929 |  |
| Sul | 6 | 5.61 | 6.01 | 0.436 | 4 | 4.18 | 3.19 | 0.156 |  |
| DFMO+Sul | 6 | 2.02 | 0.66 | 0.457 | 4 | 1.81 | 0.70 | 0.626 |  |

Note: p-value (vs. Control) by Sex was derived from one-way ANOVA and the interaction p-value between Treatment and Sex was derived from two-way ANOVA with the interaction terms between Treatment and Sex indicators

**Table 12: Summary of Spermine by treatment and sex for 58 week-old mice**

| **Treatment** | **N(Female)** | **Mean(Female)** | **SD(Female)** | **p-value (vs. Control)** | **N(Male)** | **Mean(Male)** | **SD(Male)** | **p-value (vs. Control)** | **Interaction p-value** |
| --- | --- | --- | --- | --- | --- | --- | --- | --- | --- |
| Control | 4 | 0.11 | 0.19 |  | 4 | 0.00 | 0.00 |  | 0.962 |
| DFMO | 6 | 0.10 | 0.12 | 0.973 | 5 | 0.07 | 0.10 | 0.669 |  |
| Sul | 6 | 0.32 | 0.72 | 0.423 | 4 | 0.23 | 0.43 | 0.168 |  |
| DFMO+Sul | 6 | 0.02 | 0.03 | 0.711 | 4 | 0.06 | 0.11 | 0.725 |  |

Note: p-value (vs. Control) by Sex was derived from one-way ANOVA and the interaction p-value between Treatment and Sex was derived from two-way ANOVA with the interaction terms between Treatment and Sex indicators

**Table 13: Summary of Total PA by treatment and sex for 58 week-old mice**

| **Treatment** | **N(Female)** | **Mean(Female)** | **SD(Female)** | **p-value (vs. Control)** | **N(Male)** | **Mean(Male)** | **SD(Male)** | **p-value (vs. Control)** | **Interaction p-value** |
| --- | --- | --- | --- | --- | --- | --- | --- | --- | --- |
| Control | 4 | 21.33 | 6.59 |  | 4 | 19.22 | 2.84 |  | 0.99 |
| DFMO | 6 | 18.82 | 5.55 | 0.667 | 5 | 16.49 | 3.08 | 0.207 |  |
| Sul | 6 | 22.69 | 12.33 | 0.815 | 4 | 20.59 | 3.99 | 0.537 |  |
| DFMO+Sul | 6 | 20.13 | 8.68 | 0.836 | 4 | 16.20 | 2.02 | 0.188 |  |

Note: p-value (vs. Control) by Sex was derived from one-way ANOVA and the interaction p-value between Treatment and Sex was derived from two-way ANOVA with the interaction terms between Treatment and Sex indicators

**Table 14: Summary of Sulindac by treatment and sex for 58 week-old mice**

| **Treatment** | **N(Female)** | **Mean(Female)** | **SD(Female)** | **p-value (vs. Control)** | **N(Male)** | **Mean(Male)** | **SD(Male)** | **p-value (vs. Control)** | **Interaction p-value** |
| --- | --- | --- | --- | --- | --- | --- | --- | --- | --- |
| Sul | 6 | 4309.53 | 3411.94 |  | 4 | 3592.79 | 5075.91 |  | 0.523 |
| DFMO+Sul | 6 | 6473.44 | 4608.59 | 0.377 | 4 | 3247.38 | 3748.40 | 0.916 |  |

**Table 15: Summary of Sulindac Sulfone by treatment and sex for 58 week-old mice**

| **Treatment** | **N(Female)** | **Mean(Female)** | **SD(Female)** | **p-value (vs. Control)** | **N(Male)** | **Mean(Male)** | **SD(Male)** | **p-value (vs. Control)** | **Interaction p-value** |
| --- | --- | --- | --- | --- | --- | --- | --- | --- | --- |
| Sul | 6 | 10251.33 | 3089.00 |  | 4 | 11990.54 | 4782.92 |  | 0.258 |
| DFMO+Sul | 6 | 9975.72 | 3157.35 | 0.882 | 4 | 7649.15 | 4628.41 | 0.24 |  |

**Table 16: Summary of Sulindac Sulfide by treatment and sex for 58 week-old mice**

| **Treatment** | **N(Female)** | **Mean(Female)** | **SD(Female)** | **p-value (vs. Control)** | **N(Male)** | **Mean(Male)** | **SD(Male)** | **p-value (vs. Control)** | **Interaction p-value** |
| --- | --- | --- | --- | --- | --- | --- | --- | --- | --- |
| Sul | 6 | 11991.42 | 12108.76 |  | 4 | 7519.72 | 8306.09 |  | 0.135 |
| DFMO+Sul | 6 | 1792.62 | 1259.79 | 0.067 | 4 | 9432.33 | 7966.78 | 0.751 |  |

Note: p-value (vs. Control) by Sex was derived from one-way ANOVA and the interaction p-value between Treatment and Sex was derived from two-way ANOVA with the interaction terms between Treatment and Sex indicators

**TREATMENT REGIME. Survival Analysis in young mice.**

**Table 1a: Summary of survival data by treatment for 6-week-old male mice**

| Group | N | Survival | Median (days) | Mean (days) | HR  (vs. Control) | 2.5% | 97.5% | p-value |
| --- | --- | --- | --- | --- | --- | --- | --- | --- |
| Control | 3 | 0.0 | 7 | 6.67 |  |  |  |  |
| Sodium | 5 | 0.4 | 8 | 8.00 | 0.27 | 0.05 | 1.47 | 0.13 |
| Sul | 5 | 0.0 | 6 | 5.80 | 3.88 | 0.76 | 19.89 | 0.10 |
| DFMO | 5 | 0.4 | 7 | 7.80 | 0.33 | 0.06 | 1.71 | 0.19 |
| DFMO+Sul | 5 | 0.0 | 7 | 6.60 | 0.91 | 0.22 | 3.82 | 0.90 |

**Table 1b: Summary of clinical score by treatment for 6 week-old male mice**

| Group | N | Mean | SD | p-value (vs. Control) |
| --- | --- | --- | --- | --- |
| Control | 3 | 8.33 | 0.58 |  |
| Sodium | 5 | 6.00 | 3.39 | 0.19 |
| Sul | 5 | 9.40 | 1.14 | 0.54 |
| DFMO | 5 | 6.60 | 3.36 | 0.33 |
| DFMO+Sul | 5 | 9.60 | 0.89 | 0.47 |

**Table 2a: Summary of survival data by treatment for 6-week-old female mice**

| Group | N | Survival | Median (days) | Mean (days) | HR  (vs. Control) | 2.5% | 97.5% | p-value |
| --- | --- | --- | --- | --- | --- | --- | --- | --- |
| Control | 3 | 0.0 | 7 | 6.67 |  |  |  |  |
| Sodium | 5 | 0.6 | NA | 8.60 | 0.28 | 0.05 | 1.68 | 0.16 |
| Sul | 5 | 0.2 | 7 | 7.20 | 0.75 | 0.17 | 3.39 | 0.71 |
| DFMO | 5 | 0.2 | 6 | 6.80 | 1.09 | 0.24 | 4.94 | 0.91 |
| DFMO+Sul | 5 | 0.0 | 6 | 6.00 | 1.73 | 0.4 | 7.36 | 0.46 |

**Table 2b: Summary of clinical score by treatment for 6 week-old female mice**

| Group | N | Mean | SD | p-value (vs. Control) |
| --- | --- | --- | --- | --- |
| Control | 3 | 8.33 | 0.58 |  |
| Sodium | 5 | 5.00 | 3.39 | 0.1 |
| Sul | 5 | 7.80 | 2.17 | 0.79 |
| DFMO | 5 | 7.60 | 3.78 | 0.71 |
| DFMO+Sul | 5 | 8.60 | 0.89 | 0.89 |

**Table 3a: Summary of survival data by treatment for 7-week-old female mice**

| **Group** | **N** | **Survival** | **Median (days)** | **Mean (days)** | **HR**  **(vs. Control)** | **2.5%** | **97.5%** | **p-value** |
| --- | --- | --- | --- | --- | --- | --- | --- | --- |
| Control | 3 | 0.00 | 7.0 | 6.67 |  |  |  |  |
| MilliQ | 4 | 0.50 | 7.0 | 8.00 | 0.4 | 0.0  6 | 2.45 | 0.32 |
| NaOH | 4 | 0.25 | 6.0 | 7.00 | 0.94 | 0.18 | 4.77 | 0.94 |
| DFMO MilliQ | 4 | 0.00 | 6.0 | 5.75 | 2.83 | 0.58 | 13.73 | 0.2 |
| Sul NaOH | 4 | 0.25 | 7.5 | 7.75 | 0.65 | 0.13 | 3.35 | 0.61 |

**Table 3b: Summary of clinical score by treatment for 6-week-old male mice**

| **Group** | **N** | **Mean** | **SD** | **p-value (vs. Control)** |
| --- | --- | --- | --- | --- |
| Control | 3 | 8.33 | 0.58 |  |
| MilliQ | 4 | 4.50 | 4.65 | 0.14 |
| NaOH | 4 | 6.50 | 3.70 | 0.46 |
| DFMO MilliQ | 4 | 9.25 | 0.50 | 0.71 |
| Sul NaOH | 4 | 7.00 | 3.37 | 0.59 |

**TREATMENT REGIME. Polyamine content in young mice**

**Table 1a: Summary of plasma polyamine contents by treatment for all 6-week-old male mice**

| **Polyamines by group** | **N** | **Mean** | **SD** | **p-value (vs. not infected)** |
| --- | --- | --- | --- | --- |
| **Putrescine** |  |  |  |  |
| Not Infected | 5 | 0.59 | 0.18 |  |
| Infected: Untreated | 4 | 18.27 | 3.48 | <0.0001 |
| Infected: Sodium | 5 | 2.18 | 0.91 | 0.25 |
| Infected: DFMO | 5 | 0.65 | 0.86 | 0.97 |
| Infected: Sul | 5 | 4.85 | 3.69 | <0.01 |
| Infected: DFMO+Sul | 5 | 2.38 | 1.30 | 0.20 |
| **Spermidine** |  |  |  |  |
| Not Infected | 5 | 2.37 | 0.83 |  |
| Infected: Untreated | 4 | 1.47 | 0.63 | 0.34 |
| Infected: Sodium | 5 | 2.87 | 2.16 | 0.57 |
| Infected: DFMO | 5 | 1.98 | 1.27 | 0.66 |
| Infected: Sul | 5 | 3.84 | 1.49 | 0.10 |
| Infected: DFMO+Sul | 5 | 3.73 | 1.14 | 0.13 |
| **Spermine** |  |  |  |  |
| Not Infected | 5 | 0.28 | 0.12 |  |
| Infected: Untreated | 4 | 0.03 | 0.03 | 0.50 |
| Infected: Sodium | 5 | 0.05 | 0.11 | 0.51 |
| Infected: DFMO | 5 | 0.46 | 0.62 | 0.62 |
| Infected: Sul | 5 | 2.00 | 0.85 | <0.0001 |
| Infected: DFMO+Sul | 5 | 1.43 | 0.76 | <0.01 |
| **Total polyamines** |  |  |  |  |
| Not infected | 5 | 3.24 | 1.00 |  |
| Infected: Untreated | 4 | 19.78 | 4.10 | <0.0001 |
| Infected: Sodium | 5 | 5.10 | 2.82 | 0.33 |
| Infected: DFMO | 5 | 3.08 | 2.67 | 0.93 |
| Infected: Sul | 5 | 10.68 | 4.12 | <0.001 |
| Infected: DFMO+Sul | 5 | 7.54 | 2.23 | 0.03 |

**Table 1b: Summary of plasma polyamine contents by treatment for infected 6-week-old male mice**

| **Polyamines by group** | **N** | **Mean** | **SD** | **p-value (vs. untreated)** |
| --- | --- | --- | --- | --- |
| **Putrescine** |  |  |  |  |
| Infected: Untreated | 4 | 18.27 | 3.48 |  |
| Infected: Sodium | 5 | 2.18 | 0.91 | <0.0001 |
| Infected: DFMO | 5 | 0.65 | 0.86 | <0.0001 |
| Infected: Sul | 5 | 4.85 | 3.69 | <0.0001 |
| Infected: DFMO+Sul | 5 | 2.38 | 1.30 | <0.0001 |
| **Spermidine** |  |  |  |  |
| Infected: Untreated | 4 | 1.47 | 0.63 |  |
| Infected: Sodium | 5 | 2.87 | 2.16 | 0.17 |
| Infected: DFMO | 5 | 1.98 | 1.27 | 0.61 |
| Infected: Sul | 5 | 3.84 | 1.49 | 0.03 |
| Infected: DFMO+Sul | 5 | 3.73 | 1.14 | 0.03 |
| **Spermine** |  |  |  |  |
| Infected: Untreated | 4 | 0.03 | 0.03 |  |
| Infected: Sodium | 5 | 0.05 | 0.11 | 0.97 |
| Infected: DFMO | 5 | 0.46 | 0.62 | 0.3 |
| Infected: Sul | 5 | 2.00 | 0.85 | <0.0001 |
| Infected: DFMO+Sul | 5 | 1.43 | 0.76 | <0.01 |
| **Total polyamines** |  |  |  |  |
| Infected: Untreated | 4 | 19.78 | 4.10 |  |
| Infected: Sodium | 5 | 5.10 | 2.82 | <0.0001 |
| Infected: DFMO | 5 | 3.08 | 2.67 | <0.0001 |
| Infected: Sul | 5 | 10.68 | 4.12 | <0.001 |
| Infected: DFMO+Sul | 5 | 7.54 | 2.23 | <0.0001 |

**Table 1c: Summary of Sulindac metabolites by treatment for infected 6-week-old male mice**

| **Metabolites by group** | **N** | **Mean** | **SD** | **p-value (vs. Sul)** |
| --- | --- | --- | --- | --- |
| **Sulindac** |  |  |  |  |
| Infected: Sul | 5 | 11105.68 | 10442.08 |  |
| Infected: DFMO+Sul | 5 | 12879.12 | 10466.35 | 0.8 |
| **Sulindac Sulfide** |  |  |  |  |
| Infected: Sul | 5 | 4207.47 | 4363.02 |  |
| Infected: DFMO+Sul | 5 | 2039.03 | 1525.07 | 0.32 |
| **Sulindac Sulfone** |  |  |  |  |
| Infected: Sul | 5 | 17322.07 | 17829.82 |  |
| Infected: DFMO+Sul | 5 | 19098.69 | 10452.43 | 0.85 |

**Table 2a: Summary of plasma polyamine contents by treatment for all 6-week-old female mice**

| **Polyamines by group** | **N** | **Mean** | **SD** | **p-value (vs. not infected)** |
| --- | --- | --- | --- | --- |
| **Putrescine** |  |  |  |  |
| Not Infected | 4 | 13.64 | 2.24 |  |
| Infected: Untreated | 3 | 18.06 | 1.97 | 0.98 |
| Infected: Sodium | 5 | 14.69 | 0.79 | 0.99 |
| Infected: DFMO | 5 | 18.50 | 7.21 | 0.98 |
| Infected: Sul | 5 | 303.98 | 553.03 | 0.09 |
| Infected: DFMO+Sul | 5 | 36.01 | 43.45 | 0.89 |
| **Spermidine** |  |  |  |  |
| Not Infected | 4 | 4.10 | 4.32 |  |
| Infected: Untreated | 3 | 3.78 | 2.45 | 0.91 |
| Infected: Sodium | 5 | 0.04 | 0.08 | 0.09 |
| Infected: DFMO | 5 | 0.24 | 0.44 | 0.11 |
| Infected: Sul | 5 | 3.10 | 6.66 | 0.67 |
| Infected: DFMO+Sul | 5 | 0.18 | 0.24 | 0.1 |
| **Spermine** |  |  |  |  |
| Not Infected | 4 | 0.21 | 0.37 |  |
| Infected: Untreated | 3 | 0.20 | 0.29 | 1 |
| Infected: Sodium | 5 | 3.07 | 1.00 | 0.94 |
| Infected: DFMO | 5 | 4.70 | 4.68 | 0.9 |
| Infected: Sul | 5 | 61.37 | 122.56 | 0.1 |
| Infected: DFMO+Sul | 5 | 5.90 | 4.72 | 0.88 |
| **Total polyamines** |  |  |  |  |
| Not Infected | 4 | 17.95 | 6.54 |  |
| Infected: Untreated | 3 | 22.05 | 4.39 | 0.99 |
| Infected: Sodium | 5 | 17.80 | 1.58 | 1 |
| Infected: DFMO | 5 | 23.44 | 11.73 | 0.98 |
| Infected: Sul | 5 | 368.45 | 682.16 | 0.09 |
| Infected: DFMO+Sul | 5 | 42.09 | 47.44 | 0.91 |

**Table 2b: Summary of plasma polyamine contents by treatment for infected 6-week-old female mice**

| **Polyamines by group** | **N** | **Mean** | **SD** | **p-value (vs. untreated)** |
| --- | --- | --- | --- | --- |
| **Putrescine** |  |  |  |  |
| Infected: Untreated | 3 | 18.06 | 1.97 |  |
| Infected: Sodium | 5 | 14.69 | 0.79 | 0.99 |
| Infected: DFMO | 5 | 18.50 | 7.21 | 1 |
| Infected: Sul | 5 | 303.98 | 553.03 | 0.15 |
| Infected: DFMO+Sul | 5 | 36.01 | 43.45 | 0.93 |
| **Spermidine** |  |  |  |  |
| Infected: Untreated | 3 | 3.78 | 2.45 |  |
| Infected: Sodium | 5 | 0.04 | 0.08 | 0.13 |
| Infected: DFMO | 5 | 0.24 | 0.44 | 0.15 |
| Infected: Sul | 5 | 3.10 | 6.66 | 0.78 |
| Infected: DFMO+Sul | 5 | 0.18 | 0.24 | 0.15 |
| **Spermine** |  |  |  |  |
| Infected: Untreated | 3 | 0.20 | 0.29 |  |
| Infected: Sodium | 5 | 3.07 | 1.00 | 0.95 |
| Infected: DFMO | 5 | 4.70 | 4.68 | 0.92 |
| Infected: Sul | 5 | 61.37 | 122.56 | 0.16 |
| Infected: DFMO+Sul | 5 | 5.90 | 4.72 | 0.89 |
| **Total polyamines** |  |  |  |  |
| Infected: Untreated | 3 | 22.05 | 4.39 |  |
| Infected: Sodium | 5 | 17.80 | 1.58 | 0.99 |
| Infected: DFMO | 5 | 23.44 | 11.73 | 1 |
| Infected: Sul | 5 | 368.45 | 682.16 | 0.16 |
| Infected: DFMO+Sul | 5 | 42.09 | 47.44 | 0.93 |

**Table 2b: Summary of Sulindac metabolites by treatment infected 6-week-old female mice**

| **Metabolites by group** | **N** | **Mean** | **SD** | **p-value (vs. Sul)** |
| --- | --- | --- | --- | --- |
| **Sulindac** |  |  |  |  |
| Infected: Sul | 5 | 5152.45 | 3199.21 |  |
| Infected: DFMO+Sul | 5 | 7625.51 | 6257.21 | 0.45 |
| **Sulindac Sulfide** |  |  |  |  |
| Infected: Sul | 5 | 1784.43 | 734.56 |  |
| Infected: DFMO+Sul | 5 | 8828.69 | 7984.65 | 0.09 |
| **Sulindac Sulfone** |  |  |  |  |
| Infected: Sul | 5 | 5546.65 | 2929.11 |  |
| Infected: DFMO+Sul | 5 | 7473.67 | 7494.03 | 0.61 |

**TREATMENT REGIME. Survival analysis in aged mice**

**Table 1: Summary of survival data by treatment for 58 week-old female mice**

| Group | N | Survival | Median (days) | Mean (days) | HR  (vs. Control) | 2.5% | 97.5% | p-value |
| --- | --- | --- | --- | --- | --- | --- | --- | --- |
| Control | 4 | 0.5 | 11 | 11.50 |  |  |  |  |
| DFMO | 5 | 0.6 | NA | 11.60 | 0.902 | 0.127 | 6.409 | 0.918 |
| Sul | 5 | 0.4 | 10 | 11.00 | 1.344 | 0.223 | 8.112 | 0.747 |
| DFMO+Sul | 5 | 0.0 | 9 | 9.00 | 4.125 | 0.718 | 23.706 | 0.112 |

**Table 2: Summary of clinical score by treatment for 58 week-old female mice**

| **Group** | **N** |  | **Mean** | **SD** | **p-value (vs. Control)** |
| --- | --- | --- | --- | --- | --- |
| Control | 4 |  | 5.0 | 1.83 |  |
| DFMO | 5 |  | 4.6 | 4.16 | 0.85 |
| Sul | 5 |  | 7.0 | 3.94 | 0.351 |
| DFMO+Sul | 5 |  | 8.8 | 0.84 | 0.087 |

**Table 3: Summary of survival data by treatment for 58 week-old male mice**

| **Group** | **N** | **Survival** | **Median (days)** | **Mean (days)** | **HR (vs. Control)** | **2.5%** | **97.5%** | **p-value** |
| --- | --- | --- | --- | --- | --- | --- | --- | --- |
| Control | 4 | 0.25 | 9 | 9.75 |  |  |  |  |
| DFMO | 4 | 0.50 | 9 | 11.00 | 0.593 | 0.099 | 3.559 | 0.567 |
| Sul | 4 | 0.50 | 9 | 11.00 | 0.593 | 0.099 | 3.559 | 0.567 |
| DFMO+Sul | 4 | 0.75 | NA | 12.75 | 0.231 | 0.024 | 2.235 | 0.206 |

**Table 4: Summary of clinical score by treatment for 58-week-old male mice**

| Group | N | Mean | SD | p-value (vs. Control) |
| --- | --- | --- | --- | --- |
| Control | 4 | 8.75 | 4.03 |  |
| DFMO | 4 | 6.25 | 4.92 | 0.398 |
| Sul | 4 | 5.75 | 3.78 | 0.313 |
| DFMO+Sul | 4 | 4.25 | 3.20 | 0.14 |

**TREATMENT REGIME. Polyamine content in aged mice**

**Table 3a: Summary of plasma polyamine contents by treatment for infected 58 week-old male mice**

| **Polyamines by group** | **N** | **Mean** | **SD** | **p-value (vs. untreated)** |
| --- | --- | --- | --- | --- |
| **Putrescine** |  |  |  |  |
| Infected: Untreated | 3 | 14.11 | 1.81 |  |
| Infected: DFMO | 4 | 12.96 | 2.51 | 0.63 |
| Infected: Sul | 4 | 20.76 | 4.93 | 0.02 |
| Infected: DFMO+Sul | 4 | 15.54 | 1.38 | 0.56 |
| **Spermidine** |  |  |  |  |
| Infected: Untreated | 3 | 1.11 | 0.07 |  |
| Infected: DFMO | 4 | 1.36 | 0.37 | 0.86 |
| Infected: Sul | 4 | 2.24 | 0.77 | 0.43 |
| Infected: DFMO+Sul | 4 | 3.51 | 3.34 | 0.11 |
| **Spermine** |  |  |  |  |
| Infected: Untreated | 3 | 0.04 | 0.04 |  |
| Infected: DFMO | 4 | 0.10 | 0.03 | 0.16 |
| Infected: Sul | 4 | 0.11 | 0.06 | 0.11 |
| Infected: DFMO+Sul | 4 | 0.06 | 0.07 | 0.62 |
| **Total PA** |  |  |  |  |
| Infected: Untreated | 3 | 15.26 | 1.91 |  |
| Infected: DFMO | 4 | 14.42 | 2.81 | 0.73 |
| Infected: Sul | 4 | 23.11 | 4.40 | <0.01 |
| Infected: DFMO+Sul | 4 | 19.11 | 2.31 | 0.13 |

**Table 3c: Summary of Sulindac metabolites by treatment for infected 58 week-old male mice**

| **Metabolites by group** | **N** | **Mean** | **SD** | **p-value (vs. Sul)** |
| --- | --- | --- | --- | --- |
| **Sulindac** |  |  |  |  |
| DFMO+Sul | 4 | 804.55 | 456.54 |  |
| Sul | 4 | 2006.93 | 2103.03 | 0.31 |
| **Sulindac Sulfide** |  |  |  |  |
| DFMO+Sul | 4 | 845.65 | 575.51 |  |
| Sul | 4 | 2036.49 | 2137.77 | 0.32 |
| **Sulindac Sulfone** |  |  |  |  |
| DFMO+Sul | 4 | 10204.07 | 6697.78 |  |
| Sul | 4 | 12103.51 | 6579.06 | 0.7 |

**Table 4a: Summary of plasma polyamine contents by treatment for infected 58 week-old female mice**

| **Polyamines by group** | **N** | **Mean** | **SD** | **p-value (vs. untreated)** |
| --- | --- | --- | --- | --- |
| **Putrescine** |  |  |  |  |
| Infected: Untreated | 4 | 12.05 | 0.71 |  |
| Infected: DFMO | 5 | 12.79 | 4.04 | 0.75 |
| Infected: Sul | 5 | 12.14 | 3.91 | 0.97 |
| Infected: DFMO+Sul | 5 | 13.63 | 3.07 | 0.49 |
| **Spermidine** |  |  |  |  |
| Infected: Untreated | 4 | 1.42 | 1.00 |  |
| Infected: DFMO | 5 | 2.01 | 1.54 | 0.59 |
| Infected: Sul | 5 | 3.02 | 2.42 | 0.15 |
| Infected: DFMO+Sul | 5 | 1.58 | 0.65 | 0.88 |
| **Spermine** |  |  |  |  |
| Infected: Untreated | 4 | 0.00 | 0.00 |  |
| Infected: DFMO | 5 | 0.12 | 0.15 | 0.17 |
| Infected: Sul | 5 | 0.06 | 0.14 | 0.48 |
| Infected: DFMO+Sul | 5 | 0.08 | 0.14 | 0.38 |
| **Total polyamines** |  |  |  |  |
| Infected: Untreated | 4 | 13.47 | 0.46 |  |
| Infected: DFMO | 5 | 14.92 | 5.65 | 0.65 |
| Infected: Sul | 5 | 15.22 | 5.67 | 0.58 |
| Infected: DFMO+Sul | 5 | 15.29 | 3.78 | 0.56 |

**Table 4b: Summary of Sulindac metabolites by treatment for infected 58-week-old female mice**

| **Metabolites by group** | **N** | **Mean** | **SD** | **p-value (vs. Sul)** |
| --- | --- | --- | --- | --- |
| **Sulindac** |  |  |  |  |
| DFMO+Sul | 5 | 259.56 | 229.60 |  |
| Sul | 5 | 2842.22 | 3854.47 | 0.17 |
| **Sulindac Sulfide** |  |  |  |  |
| DFMO+Sul | 5 | 232.88 | 243.81 |  |
| Sul | 5 | 2129.56 | 2562.26 | 0.14 |
| **Sulindac Sulfone** |  |  |  |  |
| DFMO+Sul | 5 | 9176.47 | 6327.38 |  |
| Sul | 5 | 20045.75 | 11026.01 | 0.09 |
